# A model of persistent post SARS-CoV-2 induced lung disease for target identification and testing of therapeutic strategies

**DOI:** 10.1101/2022.02.15.480515

**Authors:** Kenneth H. Dinnon, Sarah R. Leist, Kenichi Okuda, Hong Dang, Ethan J. Fritch, Kendra L. Gully, Gabriela De la Cruz, Mia D. Evangelista, Takanori Asakura, Rodney C. Gilmore, Padraig Hawkins, Satoko Nakano, Ande West, Alexandra Schäfer, Lisa E. Gralinski, Jamie L. Everman, Satria P. Sajuthi, Mark R. Zweigart, Stephanie Dong, Jennifer McBride, Michelle R. Cooley, Jesse B. Hines, Miriya K. Love, Steve D. Groshong, Alison VanSchoiack, Stefan J. Phelan, Yan Liang, Tyler Hether, Michael Leon, Ross E. Zumwalt, Lisa M. Barton, Eric J. Duval, Sanjay Mukhopadhyay, Edana Stroberg, Alain Borczuk, Leigh B. Thorne, Muthu K. Sakthivel, Yueh Z. Lee, James S. Hagood, Jason R. Mock, Max A. Seibold, Wanda K. O’Neal, Stephanie A. Montgomery, Richard C. Boucher, Ralph S. Baric

**Affiliations:** Department of Microbiology & Immunology, University of North Carolina at Chapel Hill, Chapel Hill, North Carolina, USA; Department of Epidemiology, University of North Carolina at Chapel Hill, Chapel Hill, North Carolina, USA; Marsico Lung Institute, University of North Carolina at Chapel Hill, Chapel Hill, North Carolina, USA; Lineberger Comprehensive Cancer Center, University of North Carolina at Chapel Hill, Chapel Hill, North Carolina, USA; Center for Genes, Environment, and Health, National Jewish Health, Denver, Colorado, USA; Golden Point Scientific Laboratories, Hoover, Alabama, USA; Division of Pathology, Department of Medicine, National Jewish Health, Denver, Colorado, USA; NanoString Technologies, Seattle, Washington, USA; Department of Pathology and Laboratory Medicine, Mayo Clinic, Rochester, Minnesota, USA; Office of the Chief Medical Examiner, Oklahoma City, Oklahoma, USA; Department of Pathology, Cleveland Clinic, Cleveland, Ohio, USA; Weill Cornell Medicine, New York, New York, USA; Department of Pathology and Laboratory Medicine, University of North Carolina at Chapel Hill, Chapel Hill, North Carolina, USA; Department of Radiology, University of North Carolina at Chapel Hill, North Carolina, USA; Biomedical Research Imaging Center, University of North Carolina at Chapel Hill, Chapel Hill, North Carolina, USA; Department of Pediatrics, Pulmonology Division and Program for Rare and Interstitial Lung Disease, University of North Carolina at Chapel Hill, Chapel Hill, North Carolina, USA; Division of Pulmonary Diseases and Critical Care Medicine, University of North Carolina at Chapel Hill, Chapel Hill, North Carolina, USA; Department of Pediatrics, National Jewish Health, Denver, Colorado, USA; Division of Pulmonary Sciences and Critical Care Medicine, Department of Medicine, University of Colorado-Denver, Denver, Colorado, USA; Rapidly Emerging Antiviral Drug Discovery Initiative, University of North Carolina at Chapel Hill, Chapel Hill, North Carolina, USA

## Abstract

COVID-19 survivors develop post-acute sequelae of SARS-CoV-2 (PASC), but the mechanistic basis of PASC-associated lung abnormalities suffers from a lack of longitudinal samples. Mouse-adapted SARS-CoV-2 MA10 produces an acute respiratory distress syndrome (ARDS) in mice similar to humans. To investigate PASC pathogenesis, studies of MA10-infected mice were extended from acute disease through clinical recovery. At 15-120 days post-virus clearance, histologic evaluation identified subpleural lesions containing collagen, proliferative fibroblasts, and chronic inflammation with tertiary lymphoid structures. Longitudinal spatial transcriptional profiling identified global reparative and fibrotic pathways dysregulated in diseased regions, similar to human COVID-19. Populations of alveolar intermediate cells, coupled with focal upregulation of pro-fibrotic markers, were identified in persistently diseased regions. Early intervention with antiviral EIDD-2801 reduced chronic disease, and early anti-fibrotic agent (nintedanib) intervention modified early disease severity. This murine model provides opportunities to identify pathways associated with persistent SARS-CoV-2 pulmonary disease and test countermeasures to ameliorate PASC.

## Introduction

The ongoing COVID-19 pandemic is caused by the severe acute respiratory syndrome coronavirus 2 (SARS-CoV-2) (*1, 2*). New antivirals, antibody therapies, vaccinations, and improved critical care strategies have diminished acute fatality rates (*3*). However, ∼40% of symptomatic and asymptomatic COVID-19 survivors develop post-acute sequelae, termed PASC or ‘long-COVID’, with features that include dyspnea, fatigue, chest pain, cognitive decline, and chronic lung disease (*4–9*). Models are urgently needed to identify early biomarkers and countermeasures to identify and prevent PASC.

COVID-19 is generally characterized as biphasic with an acute phase dominated by active SARS-CoV-2 infection and a post-viral clearance phase dominated by host reparative and immunologic processes (*10*). Human autopsy samples highlight the lung disease manifestations in patients who succumbed to COVID-19 (*11, 12*), with broad features of chronic active pneumonia (CAP), alveolar architectural destruction, dense cellularity, and pulmonary fibrosis (PF) with myofibroblast proliferation and collagen deposition (*13–19*). Survivors of previous emerging coronavirus infections reported severe post-infectious fibrotic lung sequelae long after virus clearance, and autopsy data suggest similar late sequelae will follow SARS-CoV-2 infections (*20–26*). However, elucidating the pathogenesis of post-SARS-CoV-2 lung disease is difficult because autopsy samples describe disease at single time points and are highly heterogeneous. Moreover, mechanisms describing the development of non-viral CAP and/or PF in humans are poorly understood, providing only partial roadmaps on which to base studies of SARS-CoV-2 pathogenesis (*27*). Animal models offer novel opportunities to fill these gaps in knowledge.

SARS-CoV-2 infection models in standard laboratory mice are available that produce ARDS and phenocopy age-related acute SARS-CoV-2 disease (*28, 29*), but PASC-like disease phenotypes in the lung after virus clearance have not been reported. We characterized the spatial and temporal patterns associated with long-term (120 day) pulmonary consequences of SARS-CoV-2 MA10 infection in standard BALB/c laboratory mice (*28, 29*). Lung disease in mice surviving acute SARS-CoV-2 MA10 infection was investigated using complementary virologic, histologic, and immunologic techniques supplemented with immunohistochemistry (IHC) and CT scanning. Digital spatial profiling (DSP) and RNA *in situ* hybridization (ISH) were utilized to identify transcriptional profiles during acute and chronic disease phases to characterize tissue damage and repair in mice and humans. Countermeasures to prevent lung disease sequelae for SARS-CoV-2 infection were investigated.

## Results

### SARS-CoV-2 MA10 infection produces chronic pulmonary disease

PASC outcomes were investigated in young (10-week-old) and more susceptible aged (1-year-old) mice through 120 days post infection (dpi) (*29*). To induce severe acute disease without excessive mortality, 1-year-old female BALB/c mice were inoculated intranasally with 10^3^ PFU of mouse-adapted SARS-CoV-2 MA10 (*29*). Young mice received 10^4^ PFU to achieve similar acute severe disease and lung titers (∼10^7^ PFU) at 2 dpi. Mice were necropsied at 2, 7, 15, 30, 60 and 120 dpi to measure lung viral titers and collect lungs for histopathology.

Replicating previous findings (*29*), acute infection in 1-year-old mice resulted in rapid and significant decreases in body weight and 25% mortality over 7 days compared to controls (**Fig. 1A, B**). Surviving aged mice cleared culturable infection by 15 dpi, restored lung function by 15 dpi, and recovered body weight by 30/60 dpi (100% starting weight) (**Fig. 1C-F**).

**Fig. 1:**
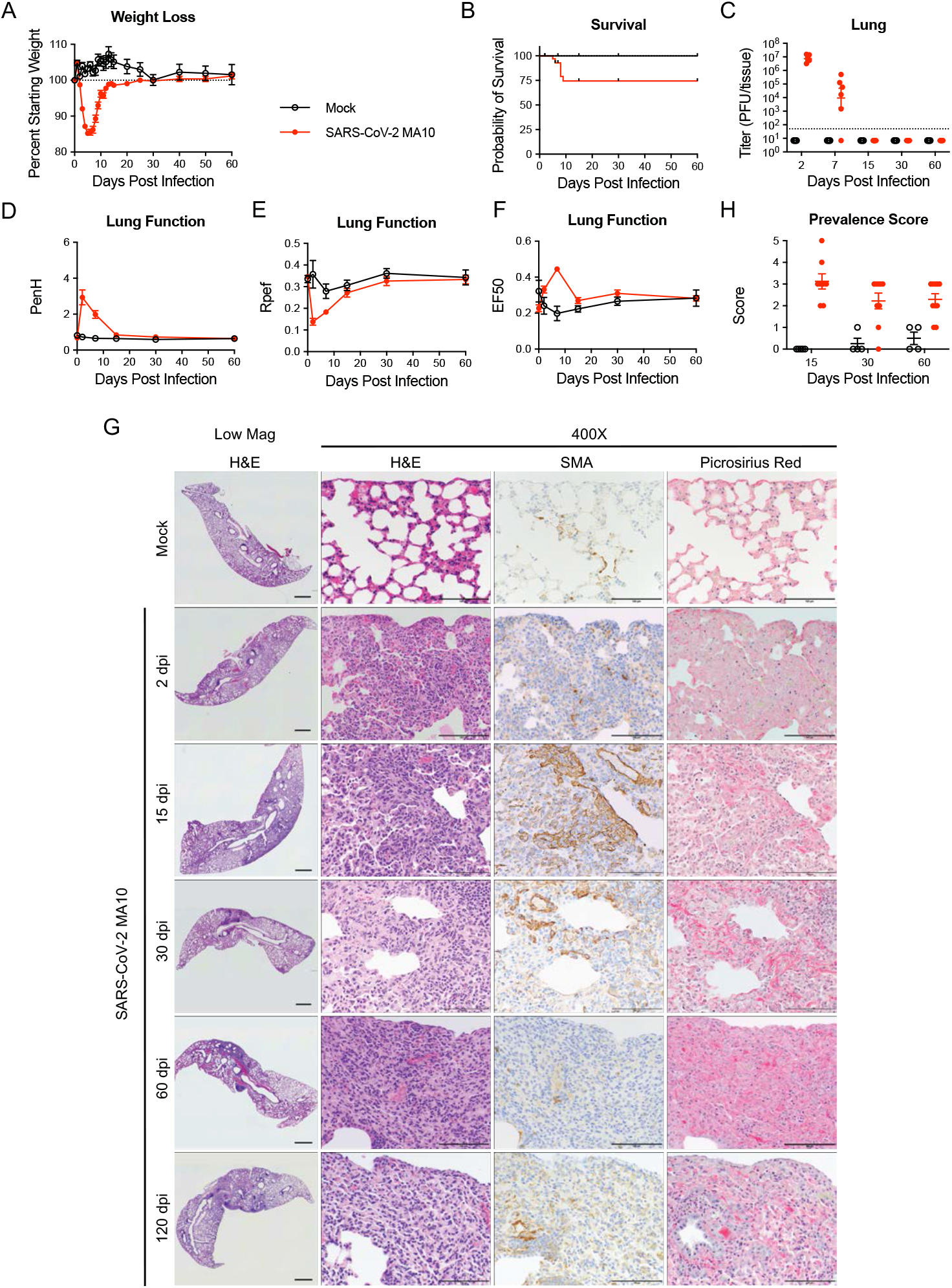
SARS-CoV-2 MA10 infection causes lung damage in aged surviving mice. 1-year-old female BALB/c mice were infected with 10^3^ PFU SARS-CoV-2 MA10 (n=74) or PBS (n=24) and monitored for **(A)** percent starting weight and **(B)** survival. **(C)** Log transformed infectious virus lung titers were assayed at indicated time points. Dotted line represents limit of detection. Undetected samples are plotted at half the limit of detection. **(D-F)** Lung function was assessed by whole body plethysmography for **(D)** PenH, **(E)** Rpef, and **(F)** EF50. **(G)** Histopathological analysis of lungs at indicated time points. H&E: hematoxylin and eosin. SMA: immunohistochemistry for α-smooth muscle actin. Picrosirius Red staining highlights collagen fibers. Image scale bars represents 1000 μm for low magnification and 100 μm for 400X images. **(H)** Disease incidence scoring at indicated time points: 0 = normal; 0 = 0% of total area of examined section, 1 = < 5%; 2 = 6-10%; 3 = 11-50%; 4 = 51-95%; 5 = > 95%. Graphs represent individuals necropsied at each timepoint (C, H), with the average value for each treatment and error bars representing standard error of the mean (A-H).

Features of acute (2-7 dpi) lung injury following SARS-CoV-2 MA10 infection in 1-year-old mice included heterogeneous inflammation and alveolar damage with consolidation, edema, fibrin and protein exudates, and occasional hyaline membranes (**Fig. 1G**) (*29*). By 15 through 120 dpi, a high incidence of histologically heterogeneous lung disease was observed (**Fig. 1G-H**). Notably, the distribution of diseased areas remained relatively constant over the 15 to 120 dpi interval, suggesting disease developed focally early and persisted. Diseased regions were often subpleurally oriented and characterized by hypercellularity with immune cell accumulation (often containing tertiary lymphoid structures), abundant smooth muscle actin (SMA) positive fibroblasts (myofibroblasts), and collagen deposition, characteristic of CAP and PF. Micro-CT scanning of 15 and 30 dpi 1-year-old mice identified dense subpleural opacities (**Fig. S1A**), and lack of honeycombing, similar to the mouse histologic lesions (**Fig. 1G-H**) and human fibrotic lung disease (*30, 31*).

Chronic manifestations were not limited to susceptible 1-year-old mice. MA10 infection (10^4^ PFU) in 10-week mice caused acute weight loss (**Fig. S2A**), 25% mortality (**Fig. S2B**), and transient pulmonary dysfunction (**Fig. S2D-E**). However, young mice cleared infectious virus earlier than old mice, by 7 dpi (**Fig. S2C**). Young mice exhibited subpleural lesions similar to old mice at 15 and 30 dpi, but the severity of disease usually diminished over 120 dpi, suggesting young mice may have a higher capacity for repair (**Fig. S2G-H**).

Cytokine analysis of lung homogenate and serum samples from both age groups revealed robust cytokine responses to infection (**Fig. S3A-B,** Supplemental Tables 1, 2). Lung cytokine responses were generally more pronounced at 2 dpi in young mice who received higher inocula. However, old mice exhibited more sustained responses post 7 dpi (**Fig. S3A**). Notably, ENA-78, M-CSF, IL-19, and Il-33, which enhance pro-fibrogenic type 2 cytokine production in a macrophage-dependent manner (*32*), remained persistently elevated in lungs to 30 dpi in older but not younger mice. In serum, a similar pattern of more robust cytokine response in young versus old mice 2 dpi was observed (**Fig. S3B**). Antiviral interferons (IFN-*α*/IFN-*λ*1) were highly expressed at 2 dpi and returned to baseline by 7 dpi at both ages (**Fig. 1C**). The more robust acute lung and plasma cytokine responses in younger versus older mice were associated with more rapid younger mouse viral clearance (by 7 dpi) (**Fig. S2C, S3**). The persistently elevated lung cytokine responses in older mice after 7 dpi may reflect delayed virus clearance and/or defective reparative capacity.

### SARS-CoV-2 MA10 infection produces acute and chronic inflammation

Immunoinflammatory responses to SARS-CoV-2 MA10 infection/injury included recruitment of macrophages, T cells, and B cells (**Fig. S4**) (*33*). Lymphoid aggregates identified in dense cellular regions at 15-120 dpi consisted of a spectrum of lymphocyte subsets, including CD4^+^, CD8^+^ T cells, and B cells (**Fig. S4A-B**). Immunohistochemistry quantitated the kinetics of CD4^+^ and CD8^+^ T cells (**Fig. S4C-D).**

Increased CD4^+^ cells appeared as early as 2 dpi, peaked at 7-15 dpi, and persisted through 120 dpi (**Fig. S4A**). CD8^+^ cell accumulation peaked at 15 dpi and remained at lower levels through 120 dpi (**Fig. S4A, D**). B220^+^ B cell accumulation was observed at 7 dpi and thereafter. CD68^+^ macrophages were increased at 7 dpi and remained elevated at 120 dpi in dense cellular regions, while iNOS^+^ M1 and Arginase^+^ M2 macrophages peaked at 2 and 7 dpi, respectively, and remained elevated at lower levels thereafter, suggesting involvement of multiple subsets of macrophages in inflammatory and reparative process with different kinetics.

Flow cytometry at 30 dpi revealed that total cells, CD45^+^ immune, and CD31^+^ endothelial cells were increased (**Fig. S4E, F**), consistent with IHC (**Fig. 4A-B**). CD4^+^ T cells and CD19^+^ B cells were significantly increased in infected mice, while CD8^+^ T cells trended higher (**Fig. S4G**), consistent with prolonged inflammatory immune responses in pulmonary fibrotic diseases (*34*). Within the monocyte/macrophage lineage, interstitial macrophages were elevated in infected mice at 30 dpi (**Fig. S4H**), consistent with a documented role that macrophages play in lung remodeling in pulmonary fibrosis (*35*).

### Spatial and temporal alteration in host transcriptional profiles in response to SARS-CoV-2 infection

GeoMx DSP was employed to interrogate viral and mouse transcripts in pulmonary lesions from a subset of mock versus infected 1-year-old mice at 2, 15, and 30 dpi (**Fig. 2A**). Since SARS-CoV-2 MA10 primarily infects alveolar AT2 cells and terminal bronchiolar secretory club cells (*29*), we focused on these two regions. At 2 dpi, alveolar regions of interest (ROIs) were selected based on the presence of SARS-CoV-2 MA10 RNA positive cells. Bronchiolar ROIs at 2 dpi were selected to represent a range of SARS-CoV-2 MA10 infection. At later time points (15, 30 dpi), the heterogeneity of alveolar lung infection/responses was sampled by obtaining ROIs from morphologically “diseased” regions with hypercellularity versus morphologically “intact” regions. All distal airways appeared normal at 15 and 30 dpi with ROIs defined as “intact”. Following data quality control/normalization, 60 alveolar and 36 bronchiolar epithelial ROIs from SARS-CoV-2 MA10-infected or mock mice were sampled at acute (2 dpi) and late (15 and 30 dpi) time points (**Fig. 2B-C, Supplemental Table 3**). Quantification of viral RNAs demonstrated clearance of viral RNAs from intact and diseased alveolar ROIs by 15 dpi (**Fig. 2D**), concordant with clearance of infectious virus (**Fig. 1C**). Normalized viral RNA counts (see Methods) trended higher in the distal airways compared with alveoli at 2 dpi and returned to baseline by 15 dpi (**Fig. 2D**).

**Fig. 2:**
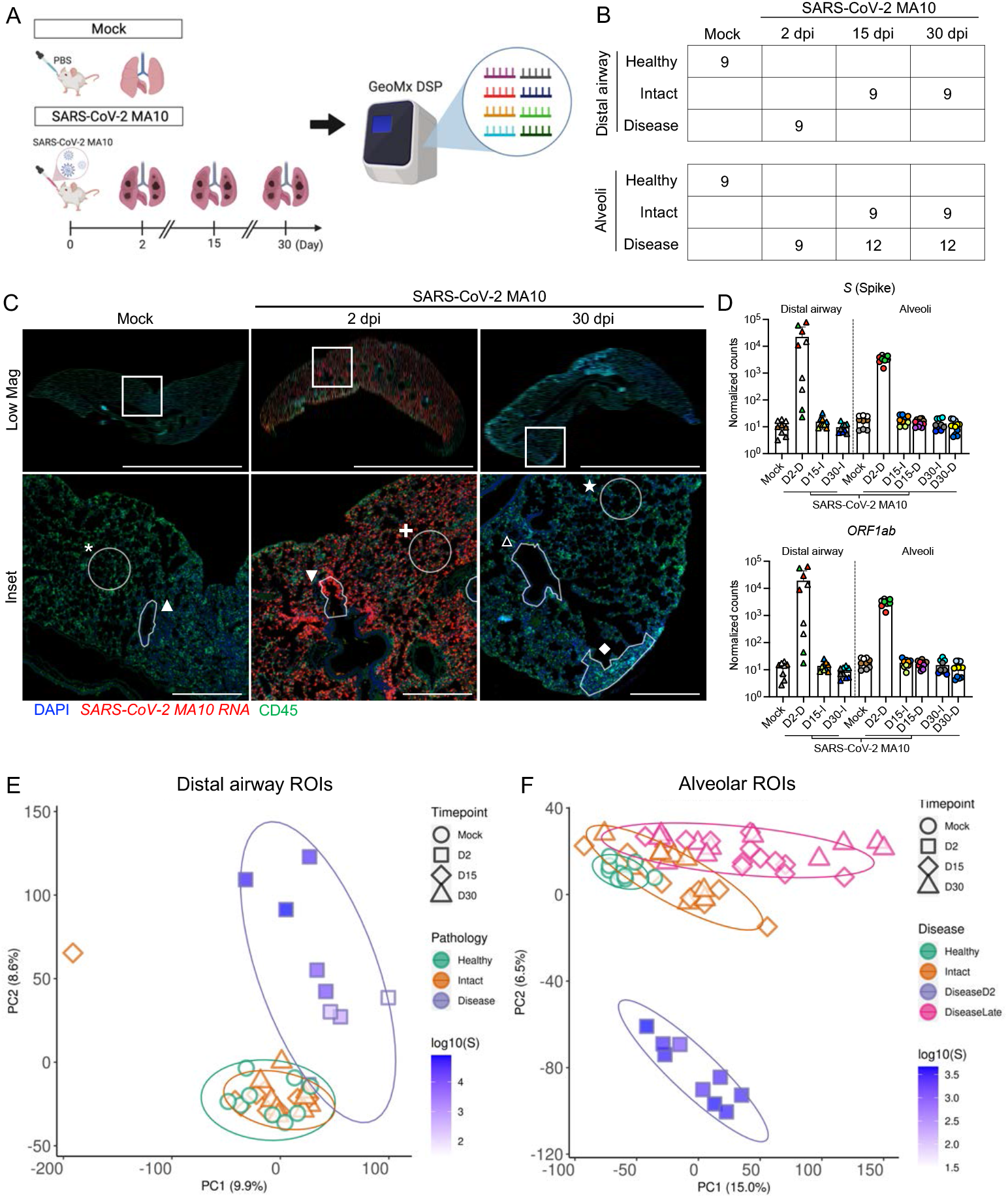
Transcriptional digital spatial profiling reveals unique signatures in diseased tissue compartments. **(A)** Experimental setup for GeoMx digital spatial profiling (DSP). **(B)** A table summarizing numbers of regions of interest (ROIs) from each tissue compartment, disease state, and time point. Each time point includes 3 independent mouse samples. **(C)** Example of ROI selections from mock, 2 dpi, and 30 dpi post SARS-CoV-2 MA10 lungs. Scale Bars = 5 mm for low magnification images and 500 μm for insets. **(D)** DSP Q3 normalized counts of SARS-CoV-2 MA10 Spike (*S*) and *ORF1ab* expression in mock, infected diseased (D), or intact (I) ROIs. Graphs represent all ROIs selected with each unique color and symbol representing one animal, bars represent average value of each group **(D)**. **(E-F)** Principal component analysis (PCA) plot of distal airway **(E)** and alveolar **(F)** ROIs.

Principal component analysis (PCA) of expressed genes identified time, region, and virus-dependent effects (**Fig. 2E, F**). High virus transcript positive regions at 2 dpi clustered away from mock in both distal airway and alveolar regions. Further, the alveolar ROIs selected from diseased regions of infected mice at 15/30 dpi separated from mock, suggesting persistent alterations of host transcriptomes (**Fig. 2F**). In contrast, the ROIs selected from “intact” airway and alveolar regions at 15/30 dpi clustered near mock healthy ROIs, suggesting recovery (**Fig. 2E-F**).

Consistent with PCA, viral infection induced major changes in transcriptome profiles in infected mouse lungs (**Fig. 3A, B; Supplemental Tables 3, 4**). In both alveoli and bronchioles, virally infected disease ROIs at 2 dpi were characterized by a broad and robust upregulation of viral infection-induced acute inflammatory genes, represented by enrichment of interferon, IL-1, and NF-kB signaling pathways (**Fig. 3A-C, Supplemental Table 5**). Upregulated ISGs were consistent with ISGs reported in human cells after emerging CoV infection (**Fig. S5A-C; Supplemental Table 2**) (*36, 37*), suggesting common antiviral pathways are activated in human and mouse pulmonary cells. As noted in other human lung cell types after CoV infection (*38*), ISG expression patterns in airway and alveolar ROIs were not identical, with some ISGs more robustly upregulated in airway epithelium (*Ifitm1*, *Lap3*, *Epsti1*) (**Fig. S5C, D**) or alveolar ROIs (*Ifitm2*, *Batf2*, *Samhd1*) (**Fig. S5C, E**). By 15 and 30 dpi, most ISG expression levels returned to mock levels (**Fig. 1C, 2D, 3A-B, Fig. S5C**).

**Fig. 3:**
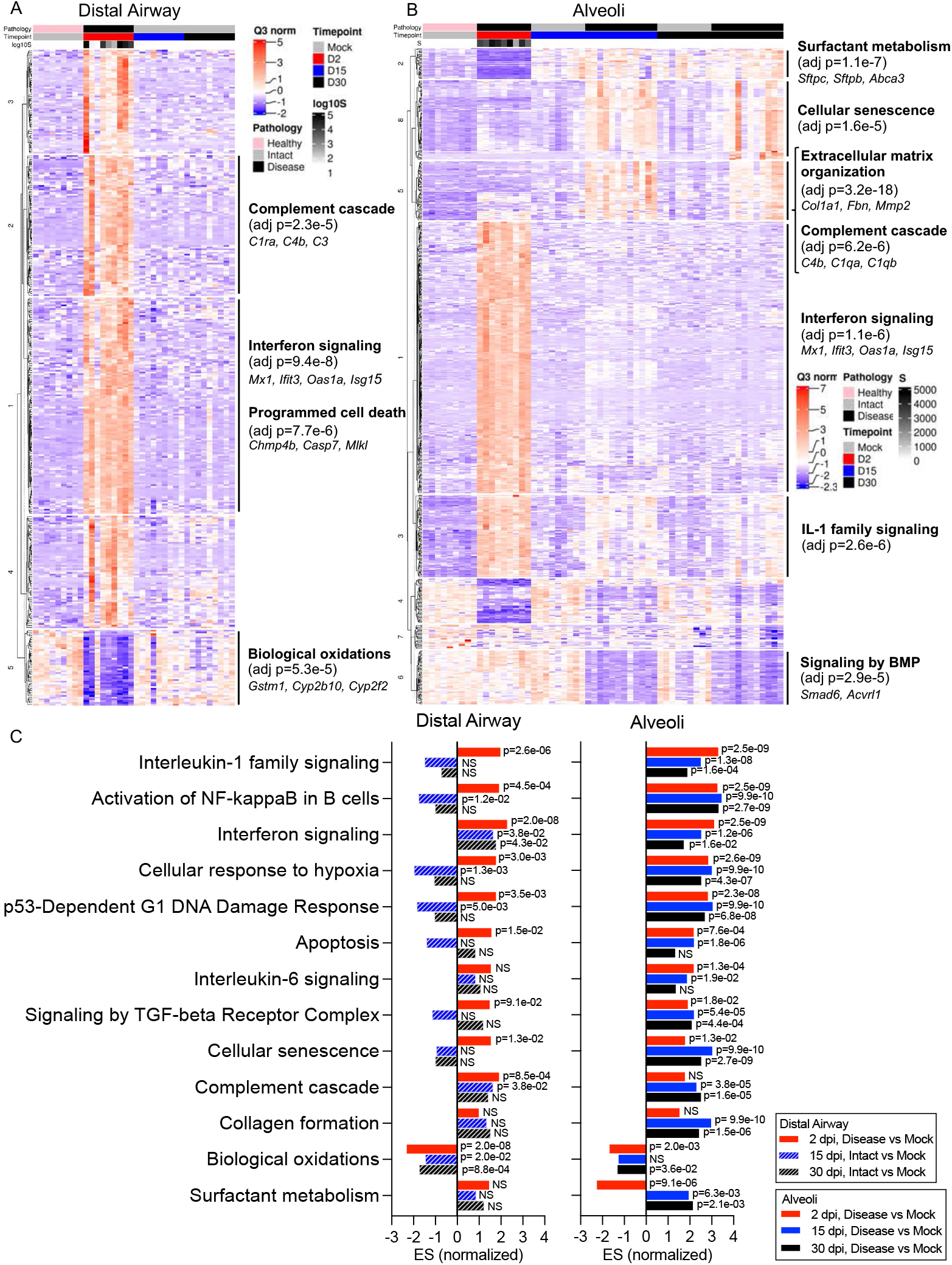
Digital spatial profiling reveals distinct transcriptional pathway changes during acute and late stages of SARS-CoV-2 disease. **(A-B)** DSP heatmaps of differentially expressed genes (DEGs) in ROIs across all time points in **(A)** distal airway and **(B)** alveolar tissue compartments. **(C)** DSP pathway enrichment analysis in distal airway and alveolar ROIs at 2, 15, and 30 dpi *vs.* mock. Statistical analyses and R packages used are detailed in methods.

DSP pathway analyses revealed downregulation of biological oxidation (bronchiolar and alveoli) and surfactant metabolism (alveoli) in infected mice at 2 dpi (**Fig. 3A-B**), associated with loss of secretory club (*Cyp2f2, Scgb1a1*, *Scgb3a2*) and AT2 (*Sftpc, Lamp3, Abca3*) cell markers (**Fig. S6A**). RNA-ISH confirmed that SARS-CoV-2 MA10 RNA was localized in *Scgb1a1*+ secretory club cells at 1 dpi and *Sftpc*+ AT2 cells at 1 dpi in bronchioles and alveoli, respectively (**Fig. S6B-C**). Significant loss of club (*Scgb1a1*) and AT2 (*Sftpc*) cell marker expression accompanied SARS-CoV-2 MA10 infection at 1-2 dpi, followed by restoration to baseline levels by 15 dpi (**Fig. S6A-E**). The early loss of *Scgb1a1* and surfactant protein genes is consistent with reported human COVID-19 autopsy data (*39*). Ciliated (*Foxj1, Dnah5, Rsph1*) and AT1 (*Ager, Hopx, Cav1*) cell markers were minimally affected by MA10 infection at any time point (**Fig. S6A-C, F**).

The transcriptomic analyses also revealed striking temporal differences in gene expression in alveolar versus bronchiolar regions (**Fig. 3A-C**). Consistent with failure of “diseased” alveolar regions to return to histologically “intact”-like states, pathway analyses at 30 dpi revealed persistently upregulated cellular senescence, hypoxia signaling, complement activation, P53 damage responses, signaling by the TGFβ receptor complex, collagen formation, and extracellular matrix organization pathways, unique to diseased alveolar regions.

The difference in post-infection histologic recovery between the bronchiolar (rapid, complete) versus alveolar regions (slow, incomplete) was notable. Because apoptosis is reported to be less inflammatory than necrotic cell death (*40*), we investigated whether apoptotic cellular responses to infection were different between the two regions (**Fig. S6G**). At 2 dpi, SARS-CoV-2 MA10-infected bronchiolar epithelial cells expressed evidence of activated apoptotic pathways (cleaved caspase-3). In contrast, alveolar regions were characterized by widespread infection but little cleaved caspase-3. These differences in apoptotic activity are consistent with reports that murine airway epithelial cells are more primed for apoptosis than alveolar epithelial cells in basal states (*41*).

### Alveolar epithelial damage and regeneration following SARS-CoV-2 infection

Recent single-cell RNA sequencing studies in acute alveolar injury mouse models have identified unique AT2 to AT1 transitional alveolar epithelial cell types following alveolar damage (*42–44*). These cells are defined variably as a Krt8+ alveolar differentiation intermediate (ADI) (*42*), damage-associated transient progenitor (DATP) (*43*), or pre-AT1 transitional state cell (PATS) (*44*) (ADI/DATP/PATS hereafter). Incomplete transition from AT2 to AT1 cells, with an accumulation of ADI/DATP/PATS cells, has also been identified in human idiopathic pulmonary fibrosis (IPF) and in COVID-19 postmortem lungs (*45, 46*), suggesting a common dysfunction in prolonged epithelial repair/disrepair. However, longitudinal characterizations of ADI/DATP/PATS cell dynamics following SARS-CoV-2 infection have not been reported.

Utilizing ADI/DATP/PATS signature genes reported from mouse acute lung injury (ALI) models (*42–44*), the SARS-CoV-2 MA10 DSP data demonstrated enrichment of ADI/DATP/PATS signatures in diseased alveolar ROIs at 2, 15, and 30 dpi (**Fig. 4A**). The ADI/DATP/PATS signature genes were categorized into three expression clusters (**Fig. 4B, Supplemental Table 2**). The first cluster (*Cdkn1a/F3/Timp1*) was enriched in diseased ROIs at 2 dpi and decreased after 15 dpi, suggesting these genes may play a role in AT2 cell trans-differentiation into ADI/DATP/PATS cells. The second cluster (*Krt8/Cxcl16/Cstb*) exhibited increased expression levels at 2 dpi through 30 dpi. The third gene cluster (*Clu*/*Eef1a1*), including a variety of ribosomal protein genes, exhibited increased expression levels at 15 dpi and later. The murine DSP gene signatures exhibited features similar to ADI/DATP/PATS signature genes identified in human COVID-19 autopsy lungs (*45*) (**Fig. S7A, Supplemental Table 2**), including p53, apoptosis, and hypoxia pathways (**Fig. 3B, C**).

To further characterize the relationships between ADI/DATP/PATS cells and disease, combined RNA-ISH and DSP analyses of reported transitional ADI/DATP/PATS cell markers (*Cdkn1a*, *Krt8*) *46*) were serially performed post infection (**Fig. 4C-D, S7B**). DSP data demonstrated that: 1) *Cdkn1a* was upregulated at 2 dpi and waned at late time points; and 2) *Krt8* was also upregulated at 2 dpi but exhibited a trend toward modestly higher expression in diseased versus intact ROIs at all points (**Fig. 4C**). While *Krt8*+/*Cdkn1a*+ RNA-ISH signals were not detectable in alveolar regions in mock mice, increased numbers of dual *Krt8+* and *Cdkn1a+* cells was observed by RNA-ISH in SARS-CoV-2-infected alveolar regions at 1-2 dpi (**Fig. 4D, S7B**), consistent with the DSP data (**Fig. 4B, C**). Notably, *Sftpc*+ AT2 cells remaining in infected alveolar regions at 1 dpi co-expressed *Krt8* and *Cdkn1a* (**Fig. S7B**), consistent with the reported AT2 to ADI/DATP/PATS transitions after ALI in mice (*42–44*). At 2 dpi, *Krt8*+/*Cdkn1a*+ cells were present and *Sftpc*+/*Krt8*+ cells were rare (**Fig. 4D, S7B**), consistent with the loss of *Sftpc* in disease ROIs at 2 dpi (**Fig. S6D, E**). At 7-15 dpi, *Sftpc* expression was restored and only occasional *Sftpc*+/*Krt8*+ cells were observed in repairing regions (**Fig. S7B**). Given the decreased viral titer (**Fig. 1C**) and restoration of *Sftpc* expression at 7-15 dpi (**Fig. S6D, G**), *Sftpc*+/*Krt8*+ cells observed in these repairing regions likely reflected *Krt8*+ ADI/DATP/PATS cells re-transitioning into mature alveolar cells. Consistent with this notion, immunohistochemistry revealed co-expression of Krt8 with both AT1 (Ager) and AT2 (Sftpc) cell markers at 30 dpi (**Fig. S7C**). However, while *Sftpc*+ AT2 cells were restored in most alveolar regions at 15-30 dpi (**Fig. 4D, S7C**), persistent *Krt8*+ and/or *Cdkn1a*+ cell clusters, coupled with muted restoration of *Sftpc*+ cells, was identified in dense cellular subpleural fibrotic alveolar regions where Col1a1 protein accumulation coexisted (**Fig. 4D**).

### Persistent inflammation and fibrosis as a chronic manifestation in SARS-CoV-2 MA10-infected mice

In diseased alveolar ROIs at 15 and 30 dpi, multiple genes involved in adaptive immune signaling and extracellular matrix deposition were highly upregulated, consistent with a wound repair/profibrotic environment (**Fig. 5A-B**). Recent human COVID-19 autopsy and transplant lung studies identified abundant interstitial pro-fibrotic monocyte-derived macrophages characterized by increased expression of *SPP1*, *MMP9*, and *CTSZ* (*45, 47, 48*). These macrophage features, coupled with upregulated extra cellular matrix remodeling (*SPARC*, *CTSK*) and macrophage-colony stimulating factor signaling genes (*CSF1*, *CSF1R*), defined a profibrotic macrophage archetype in human IPF samples (*49*). Our DSP analyses identified features associated with this profibrotic macrophage archetype in diseased alveolar ROIs at 15 and 30 dpi, including increased *Spp1*, *Sparc, and Csf1r* expression (**Fig. 5C**). RNA-ISH confirmed a persistent increase in *Spp1* expression in SARS-CoV-2 MA10-infected mice after 7 dpi (**Fig. 5D-E**). These chronic fibrotic manifestations were consistent with IHC and flow cytometry data demonstrating increased interstitial macrophage populations during chronic SARS-CoV-2 MA10 infection (**Fig. S4H**). Additionally, adaptive immune cell signatures, e.g., immunoglobulin (*Igha*, *Igkc*, *J chain*) and MHC II complex (*H2-Ea*, *H2-Eb1*, *H2-Ab1*) genes, were upregulated in diseased alveolar ROIs at 30 dpi (**Fig. 5B**), consistent with the accumulation of interstitial macrophages and CD19^+^ B cells observed by immunohistochemistry and flow cytometry (**Fig. S4A, G-H**).

**Fig. 4:**
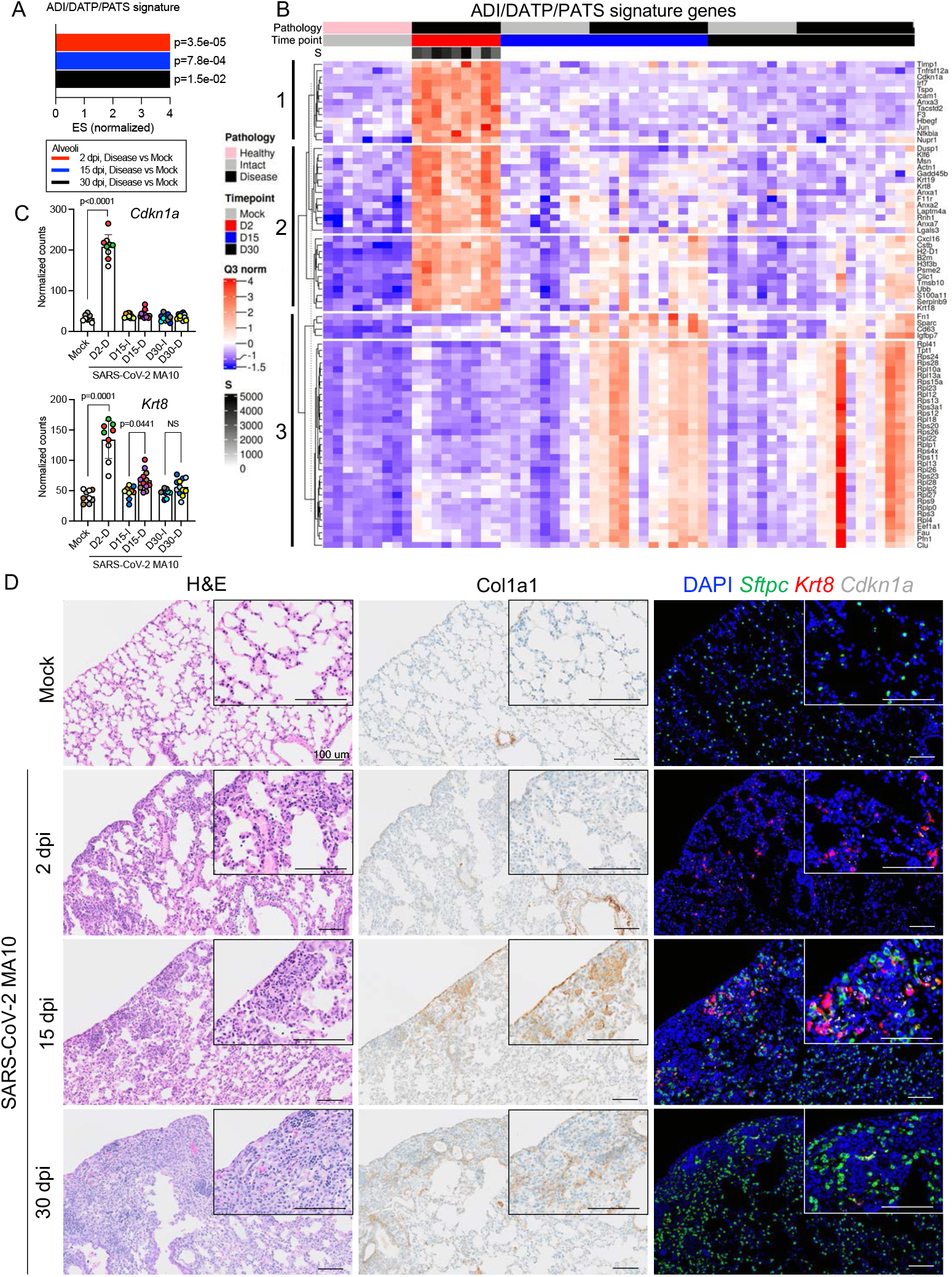
Transitional alveolar epithelial cell genes are upregulated following SARS-CoV-2 MA10 infection. **(A)** DSP pathway analysis of an ADI/DATP/PATS signature in diseased alveolar ROIs at 2, 15, and 30 dpi *vs.* mock. **(B)** DSP heatmap of reported ADI/DATP/PATS marker genes in alveolar ROIs. **(C)** DSP Q3 normalized counts of *Cdkn1a* and *Krt8* expression across alveolar ROIs. Graphs represent all ROIs selected with each unique color and symbol representing one animal, bars represent average value of each group with error bars representing standard error of the mean. The difference in DSP Q3 normalized counts for targeted genes in ROIs between each condition and time point was statistically tested using a linear mixed-effect model with condition and time point as fixed effects and replicate mice as random-effect factors. **(D)** Histopathological analysis of lungs at indicated time points. H&E: hematoxylin and eosin. Col1a1: immunohistochemistry for Col1a1. RNA-ISH for *Sftpc*, *Krt8* and *Cdkn1a*. Scale Bars = 100 μm.

**Fig. 5:**
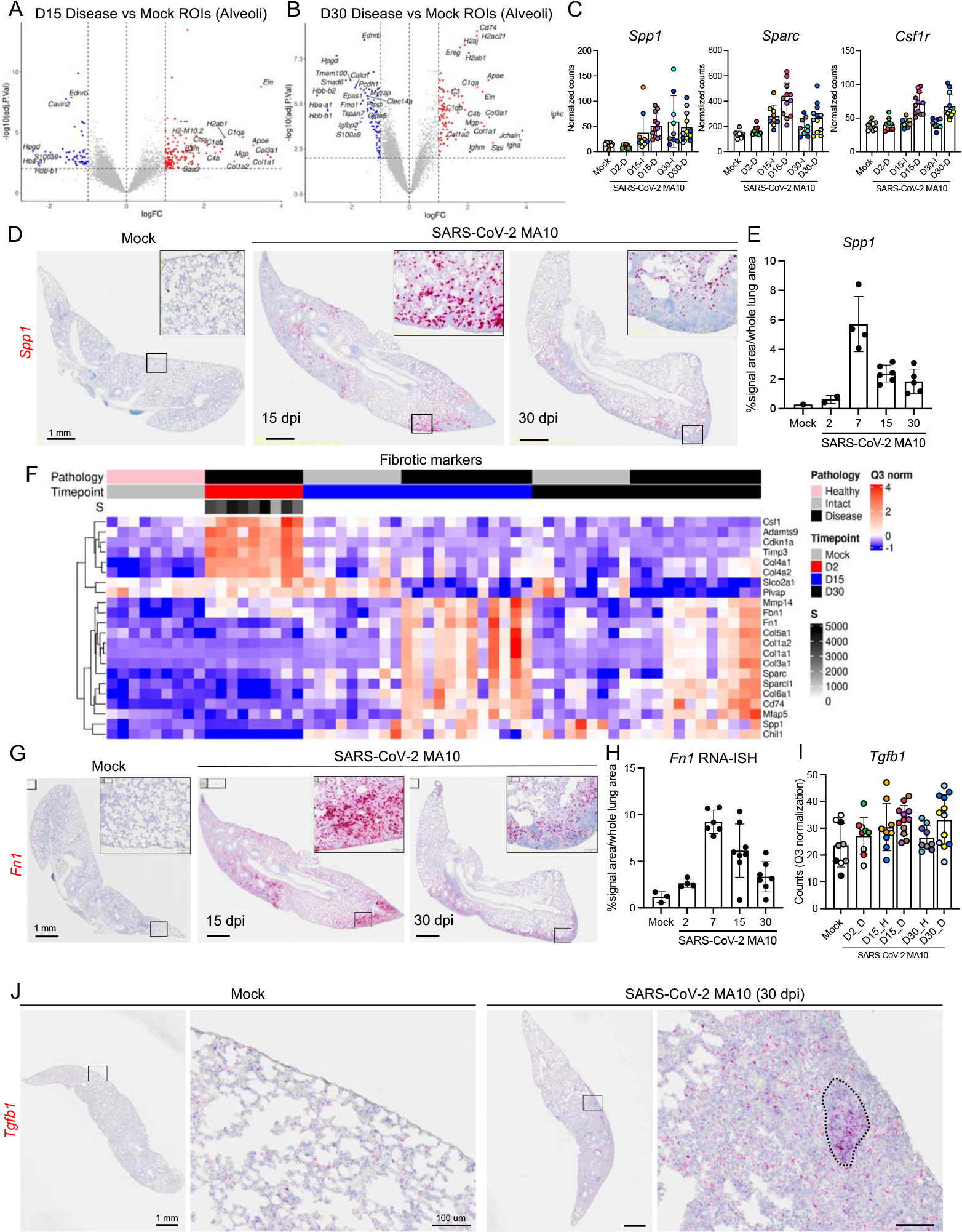
SARS-CoV-2 MA10 infection induces profibrotic gene expression at late time points. **(A-B)** Volcano plots of DSP DEGs in diseased alveolar ROIs at **(A)** 15 and **(B)** 30 dpi *vs.* mock. **(C)** DSP Q3 normalized counts of *Spp1*, *Sparc*, and *Csf1r* expression associated with profibrotic macrophage archetype. **(D-E)** *Spp1* expression by RNA-ISH **(D)** with quantification **(E)**. **(F)** DSP heatmap of selected profibrotic and fibrosis related genes in alveolar ROIs. **(G-H)** *Fn1* expression by RNA-ISH **(G)** with quantification **(H)**. **D, G**: Scale Bars = 1 mm. **(I)** DSP Q3 normalized counts of *Tgfb1*. **(J)** *Tgfb1* expression by RNA-ISH in subpleural diseased regions in a SARS-CoV-2 MA10 infected mouse at 30 dpi compared to mock. Scale Bars = 1 mm (low power) and 100 μm (high power). DSP count graphs represent all ROIs selected with each unique color and symbol representing one animal, bars represent average value of each group with error bars representing standard error of the mean **(C, I)**. RNA-ISH quantification graphs represent average value of each group with error bars representing standard error of the mean **(E, H)**.

In parallel, we characterized SARS-CoV-2 MA10-ingected mouse genes associated with human IPF (*49*). Hierarchical clustering of alveolar ROIs (**Fig. 5F**, **Supplemental Table 2**) demonstrated enrichment of extracellular matrix-related genes (*Col1a1*/*Fbn1*/*Fn1*) in mouse alveolar disease ROIs at 15 and 30 dpi (**Fig. 5A-B, F**). RNA-ISH and immunohistochemistry confirmed increased expression of Col1a1 protein and *Fn1* transcripts in the subpleural pro-fibrotic alveolar regions at 15 and 30 dpi (**Fig. 4D, 5G-H**). TGF-β is likely a central pro-fibrotic growth factor in IPF (*50*), and DSP data demonstrated an upregulated TGF-β signaling pathway (**Fig. 3C**) with trends toward *Tgfb1* upregulation in alveolar diseased versus intact ROIs at 15 and 30 dpi (**Fig. 5I**). Importantly, RNA-ISH revealed high *Tgfb1* expression in alveolar fibrotic regions, associated with lymphocyte accumulation, in SARS-CoV-2 MA10-infected mice at 30 dpi (**Fig. 5J**). These data suggest common pathways are activated in the development of IPF in humans and our mouse model of SARS-CoV-2 infection PASC.

### Mouse MA10 recapitulates features of fatal human COVID-19 lungs

We next compared mouse and published human data to a novel human COVID-19 autopsy cohort. Analyses of human COVID-19 autopsy by DSP, histology scoring, and immunohistochemistry revealed significant biological networks/processes modified by COVID-19 disease that were recapitulated in SARS-CoV-2 MA10-infected mice (**Fig. S8**). Given the small number of patients, heterogeneity of time between disease onset and death, and patient variability, pathway analyses of COVID-19 lung samples were performed rather than longitudinal/patient-based analyses. Analyses indicated: 1) significant transcriptional alteration in DSP COVID-19 ROIs separated from non-COVID ROIs indicated by PCA analysis (**Fig. S8A**); 2) histological evidence of chronic inflammation and organizing lung injury with upregulation of networks containing type I/II interferon-stimulated/IL-6-driven inflammation signatures (**Fig. S8B**); 3) upregulation of collagen/fibrotic gene signatures containing multiple human IPF genes [*COL1A1*, *COL15A1*, *FBN1*, *FN1, TNC*, consistent with mouse gene signatures; (**Fig. 5A-B, F-H)**] with significantly increased collagen and SMA protein on immunohistochemistry (**Fig. S8B-D**); 4) evidence of complement activation; an 5) evidence for altered alveolar architecture as indicated by downregulation of ATI/endothelial networks and AT2 gene markers. Note, these findings differed from mice. For example, ciliated and *TP63/MUC5AC* networks were enriched in some COVID lungs, which are consistent with histopathologic IPF features that exhibit infiltration of fibrotic alveoli with airway basal cells and “honeycombing cysts” lined by mucus producing ciliated epithelia (*50, 51*). The absence of this finding in the mouse likely reflects a dearth of basal cells in the bronchiolar region of mice and/or unknown preexisting lung disease in COVID patients (*31, 51, 52*).

### EIDD-2801 reduces chronic pulmonary lesions in mice

EIDD-2801 (molnupiravir) is an FDA approved direct-acting antiviral (DAA) that rapidly clears SARS-CoV-2 infection in mice and humans (*53, 54*). We treated infected 1-year-old female BALB/c mice with EIDD-2801 or vehicle twice daily from 12 hpi - 5 dpi post infection and followed survivors through day 30. As reported (*53*), EIDD-2801 administration reduced weight loss, mortality, virus titers, gross lung congestion, diffuse alveolar damage (DAD) and ALI during the acute phase of infection (**Fig. 6A-F**). At 30 days, profibrotic disease prevalence was significantly reduced compared to vehicle controls (**Fig. 6G-H**).

**Fig. 6:**
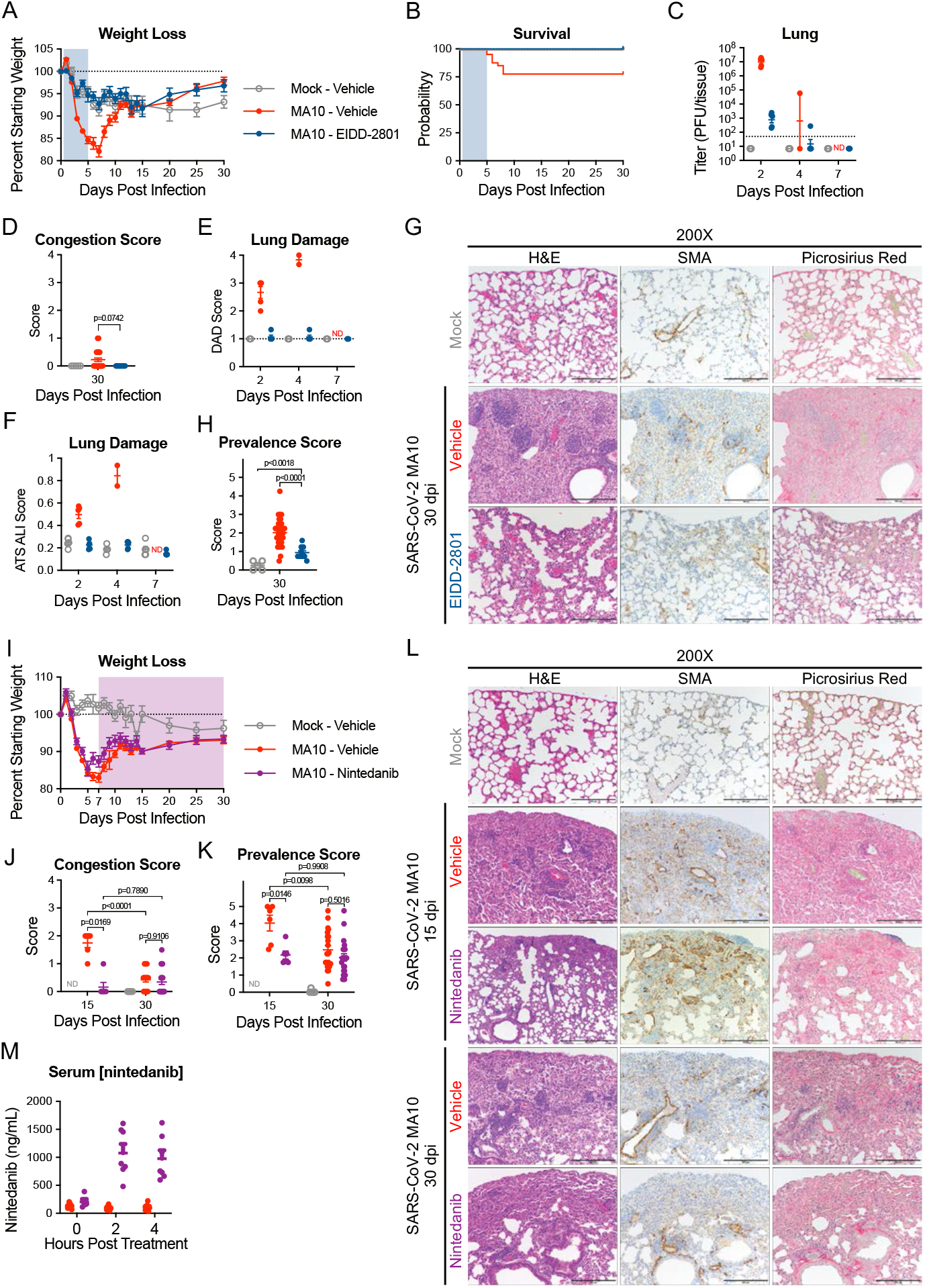
Direct acting antiviral EIDD-2801 prevents lung damage and anti-fibrotic Nintedanib reduces peak disease in SARS-CoV-2 infected aged mice. 1-year-old female BALB/c mice were infected with 10^3^ PFU of SARS-CoV-2 MA10 (n=50) or PBS (n= 5) then treated with EIDD-2801 (n= 10) (500 mg/kg BID) or vehicle (n= 45) starting at 12 hours post infection until 5 days post infection. Animals were monitored for weight loss **(A)** and survival **(B)**. Log transformed infectious virus lung titers were assayed at selected timepoints **(C)**. Dotted line indicates limit of detection and undetected samples are plotted at half the limit of detection. Pathology scores of mice as measured by lung congestion at time of harvest **(D)**, lung damage measured via evaluation of H&E staining for diffuse alveolar damage **(E)** and acute lung injury **(F)**. **(G)** Histopathological analysis of lungs at indicated time points. H&E: hematoxylin and eosin. α-SMA: immunohistochemistry for smooth muscle actin. Picrosirius Red staining highlights collagen fibers. Scale bars represents 100 μm for 200X images. **(H)** Disease incidence scoring at indicated time points: 0 = normal; 0 = 0% of total area of examined section, 1 = < 5%; 2 = 6-10%; 3 = 11-50%; 4 = 51-95%; 5 = > 95%. 1-year-old female BALB/c mice were infected with 10^3^ PFU of SARS-CoV-2 MA10 (n=90) or PBS (n=5) then treated with Nintedanib (n=45) or vehicle (n=50) starting at 7 days post infection until designated harvest date. **(I-J)** Animals were monitored for weight loss **(I)** and survival **(J)**. **(K)** Gross pathology scores of mice as measured by lung congestion at time of harvest. Disease incidence scoring at indicated time points: 0 = normal; 0 = 0% of total parenchyma, 1 = < 5%; 2 = 6-10%; 3 = 11-50%; 4 = 51-95%; 5 = > 95% **(L)** Histopathological analysis of lungs at indicated time points. H&E: hematoxylin and eosin. α-SMA: immunohistochemistry for smooth muscle actin. Picrosirius Red staining highlights collagen fibers. Image scale bars represents 100 μm for 200X images. **(M)** Serum nintedanib concentrations. Graphs represent individuals collected at each timepoint **(C-F, H-K, M)**, with the average value for each treatment and error bars representing standard error of the mean, calculated in Prism 9 **(A-F, H-K, M)**. Kruskal-Wallis **(D, H)** and two-way ANOVA **(J, K)** were performed in Prism 9 and p-values are given with comparisons on each graph.

### Nintedanib decreases peak fibrotic disease in mice

Nintedanib is an FDA approved anti-fibrotic therapeutic agent that prevents IPF progression in humans (*55, 56*). Nintedanib inhibits the tyrosine kinase PDGF, FGF, and vascular endothelial growth factor receptors and interferes with fibroblast proliferation, migration, differentiation, and secretion of extracellular matrices (*57*). Older BALB/c mice administered Nintedanib continuously from 7 dpi showed no differences in weight loss/recovery compared to vehicle treated mice through 30 dpi (**Fig. 6I**). Nintedanib treatment decreased gross tissue congestion scores, fibrotic prevalence scores, and collagen deposition, at 15 dpi compared to controls (**Fig. 6I-L**). Vehicle-treated mice exhibited reduced 30 dpi fibrotic prevalence/collagen deposition scores compared to d 15, approaching values similar to 30 dpi nintedanib-treated animals. Serum nintedanib concentrations were confirmed by UHPLC-TOF mass spectrometry to be within range previously reported in mice (*58*) (**Fig. 6M**).

## Discussion

SARS-CoV-2 infection causes acute ALI/ARDS and post-acute phase chronic lung sequelae, including CAP and PF (*59, 60*). CT scans reveal chronic COVID-19 pulmonary findings as evidenced by ground glass opacities (44%) and fibrosis (21%) after acute COVID-19 infection and fibrotic-like changes (35%) 6 months after severe human COVID-19 pneumonia (*62*). Pathology studies of COVID-19 lungs obtained at autopsy reveal similar late findings, i.e., CAP/PF (*51, 63, 64*). The SARS-CoV-2 MA10 model recapitulates these phenotypes through 120 dpi.

Currently, our understanding of PASC and COVID-19 induced CAP/PF is poor and countermeasures are limited due to the wide spectrum of potential disease pathophysiologies. Recently, a chronic (30 dpi) SARS-CoV-2 infection model was reported in immunosuppressed, humanized mice characterized by persistent virus replication and chronic inflammation with fibrotic markers, typical of rare infections seen in immunosuppressed humans who cannot clear virus (*65*). We developed a model of long-term pulmonary sequelae of SARS-CoV-2 infection that persisted after virus clearance and was more characteristic of the general patient population. In the SARS-CoV-2 MA10 model, surviving older mice cleared infection by 15 dpi but exhibited damaged pulmonary epithelia accompanied by secretion of a spectrum of pro-inflammatory/fibrotic cytokines often upregulated in fibrotic disease in humans, e.g., IL-1β, TNF-α, GM-CSF, TGF-β, IL-33, and IL-17A (**Fig. S3, 5J**) (*66*). Like humans, surviving SARS-CoV-2-infected mice by 30-120 dpi developed heterogeneous, persistent pulmonary lesions of varying severity (*67–69*) with abnormally repairing AT2 cells, interstitial macrophage and lymphoid cell accumulation, myofibroblast proliferation, and interstitial collagen deposition, particularly in subpleural regions (**Fig. 1, S1, S2**). Micro-CT detected heterogeneous subpleural opacities and fibrosis in surviving mice, similar to human studies (*70*). While most of acute cytokine production returned to normal levels by 30 dpi, DSP and RNA-ISH data revealed focally prolonged upregulation of cytokine signaling, including TGF-β, in sub-pleural fibrotic regions. Importantly, similar heterogeneous cellular and fibrotic features in subpleural regions are also evident in late stage COVID-19 patients (*71*).

SARS-CoV-2 MA10 infection caused acute loss of distal airway club cell (*Scgb1a1*) and alveoli AT2 cell (*Sftpc*) marker expression, phenotypes consistent with SARS-CoV-2 cellular tropisms in humans (*72*) (**Fig. 3, S5**). The expression levels of club/AT2 cell genes were variably restored by 15 dpi as demonstrated by DSP and RNA-ISH data (**Fig. S5**). We speculate that a key variable determining the ability of the alveolar region to repair, or not, reflects the capacity of surviving and/or residual AT2 cells to regenerate an intact alveolar epithelium. The failure of AT2 cells to replenish themselves or AT1 cells and repair alveolar surfaces in subpleural regions likely reflects the intensity of SARS-CoV-2 infection. Based on data from COVID-19 autopsy lungs, an accumulation of replication-defective/pro-inflammatory (ADI/DATP/PATS) transitional cells emerge early after SARS-CoV-2 infection and may persist, associated with persistent inflammation and failure of repair (*45, 46*). Our longitudinal mouse model data support this notion as evidenced by the observation that ADI/DATP/PATS cells were detected at 2 dpi and persisted through 30 dpi in diseased, but not morphologically intact, alveolar regions (**Fig. 4**). These ADI/DATP/PATS cells were notable for upregulation of senescence, Hif1α, and pro-inflammatory cytokines, e.g., IL-1β pathways, in keeping with low cycling rates/failure to replenish AT2/AT1 cells and a pro-inflammatory phenotype (*43*). However, as evidenced by the return of significant *Sftpc* expression by 15 dpi in intact alveolar regions, a fraction of the ADI/DATP/PATS cells likely regenerated mature *Sftpc*-expressing AT2 cells. Notably, our longitudinal studies revealed that the gene expression profiles of ADI/DATP/PATS cells are dynamic over the evolution of lung disease (**Fig. 4B, S7A**).

As reported in humans, CD4^+^/CD8^+^ lymphocyte populations increased in SARS-CoV-2-diseased areas of mouse lungs, and peripheral lymphoid aggregations were a feature of chronic disease (**Fig. 1**). These features were consistent across all analyses, including immunohistochemistry, DSP, and flow cytometry data. A notable macrophage feature, identified by DSP and flow cytometry data, was expansion of the interstitial macrophage population, consistent with human data (*47*). The subpleural regions exhibited the most striking histologic evidence of immunologic cell recruitment and activation of adaptive immune, hypoxia, fibrotic, and extracellular matrix pathways in association with ADI/DATP/PATS cells (**Fig. 3-6**).

Final clues to the etiology of the late-stage alveolar CAP/PF response emerged from comparisons to infection in bronchioles. Despite quasi-higher bronchiolar infection intensities, bronchioles repaired without evidence of organizing/fibrotic sequelae. Bronchioles may be protected from this adverse fate by tissue-specific ISG responses to control the duration/severity of infection (**Fig. S5C-E**). In this context, several ISGs, including *Ifitm1* and *Ifitm2*, exhibited clear differences in tissue specific expression and/or persistence through 30 dpi (**Fig. 2**, **S5**). Other possible relevant variables that may favor bronchiolar repair include: 1) more “controlled” cell death, i.e., apoptosis (**Fig. S6G**); 2) a less damaged basement membrane architecture; and 3) inability of club cells to enter an intermediate, ADI/DATP/PATS cell equivalent (**Fig. 4**).

Mouse models of acute and chronic viral disease are critical also for countermeasure development. Molnupiravir is one of three FDA-approved DAA that clear virus, reduce morbidity, mortality, and time to recovery (*53, 73*). Early molnupiravir treatment attenuated chronic PASC in the SARS-CoV-2 MA10 mouse model (**Fig. 6**). Although speculative, early DAA treatment may forestall chronic lung and other organ PASC manifestations. Based on preclinical studies of anti-fibrotic agents in reducing the severity of PF responses to chemical agents, we tested the concept that early intervention with an anti-fibrotic agent may reduce the severity of PF following SARS-CoV-2 infection (*57*). Nintedanib administered from 7 dpi blunted maximal fibrotic responses to virus at 15 dpi, supporting the concept that early intervention with anti-fibrotic agents may attenuate post-SARS-CoV-2 severe disease trajectories.

In summary, the SARS-CoV-2 MA10 mouse model provides novel opportunities to longitudinally study the molecular mechanisms/pathways mediating long-term COVID-19 pulmonary sequelae as relates to human PASC. The model supports high-priority research directions that include SARS-CoV-2 infection of transgenic lineage tracing reporter mice to define longitudinally the fates of infected club and AT2 cells, ADI/DATP/PATS cell transitions, mechanisms of cell death, and epithelial cell regeneration/repopulation following infection. With respect to countermeasures, ∼1 year clinical trials are required to assess therapeutic benefit for lung fibrosis, emphasizing the utility of the SARS-CoV-2 MA10 model to test rapidly agents that may counter the pulmonary CAP/PF effects of COVID-19 (*74, 75*). Thus, the murine SARS-CoV-2 MA10 model permits longitudinal selection/validation of therapeutic targets, accelerated timelines, and controlled experimental settings for testing of novel therapeutic agents.

## Material and Methods

### Ethics and biosafety

The generation of SARS-CoV-2 MA10 was approved for use under BSL3 conditions by the University of North Carolina at Chapel Hill Institutional Review Board (UNC-CH IBC) and by a Potential Pandemic Pathogen Care and Oversight committee at the National Institute of Allergy and Infectious Diseases (NIAID). All animal work was approved by Institutional Animal Care and Use Committee at University of North Carolina at Chapel Hill according to guidelines outlined by the Association for the Assessment and Accreditation of Laboratory Animal Care and the U.S. Department of Agriculture. All work was performed with approved standard operating procedures and safety conditions for SARS-CoV-2, including all virologic work was performed in a high containment BSL3 facility and personnel wore PAPR, Tyvek suits and were double gloved. Our institutional BSL3 facilities have been designed to conform to the safety requirements recommended by Biosafety in Microbiological and Biomedical Laboratories (BMBL), the U.S. Department of Health and Human Services, the Public Health Service, the Centers for Disease Control and Prevention (CDC), and the National Institutes of Health (NIH). Laboratory safety plans have been submitted, and the facility has been approved for use by the UNC Department of Environmental Health and Safety (EHS) and the CDC.

### Viruses and cells

Serial *in vivo* passaging of parental SARS-CoV-2 MA virus (*76*) in mice lead to the plaque purification of a passage 10 clonal isolate (SARS-CoV-2 MA10) (*29*). A large working stock of SARS-CoV-2 MA10 was generated by passaging the plaque purified clonal isolate sequentially on Vero E6 cells at 37°C (passage 3, SARS-CoV-2 P3). SARS-CoV-2 MA10 P3 was used for all *in vivo* experiments.

Vero E6 cells were cultured in Dulbecco’s modified Eagle’s medium (DMEM, Gibco) with the addition of 5% Fetal Clone II serum (Hyclone) and 1X antibiotic/antimycotic (Gibco). Working stock titers were determined via plaque assay by adding serially diluted virus to Vero E6 cell monolayers. After incubation, monolayers were overlayed with media containing 0.8% agarose. After 72 hours, Neutral Red dye was used to visualize plaques.

### *In vivo* infection

All BALB/c mice used in this study were purchased from Envigo (BALB/cAnNHsd; strain 047) and housed at the University of North Carolina at Chapel Hill until the start of the experiment. For intranasal infection, mice were anesthetized using a mixture of ketamine and xylazine. 10^4^ plaque forming units (PFU) or 10^3^ PFU of SARS-CoV-2 MA10 diluted in PBS were used for inoculation of young (10 week) or aged (12 months) BALB/c mice, respectively. Weight loss and morbidity were monitored daily as clinical signs of disease whereas lung function was assessed at indicated time points using whole body plethysmography (WBP; DSI Buxco respiratory solutions, DSI Inc.). Lung function data was acquired as previously described (*77*) by allowing mice to acclimate in WBP chambers for 30 min and a data acquisition time of 5 min. Data was analyzed using FinePointe software.

At indicated harvest time points, randomly assigned animals were euthanized by an overdose of isoflurane and samples for analyses of titer (caudal right lung lobe) and histopathology (left lung lobe) were collected. Animals recorded as “dead” on non-harvest days were either found dead in cage or were approaching 70% of their starting body weight which resembles the criteria for humane euthanasia defined by respective animal protocols.

Viral titers in lungs were determined by plaque assay for which caudal right lung lobes were homogenized in 1mL of PBS and glass beads, monolayers of Vero E6 cells inoculated, and 72 hours after incubation stained with Neutral Red dye for visualization of plaques.

### Disease incidence scoring

Profibrotic disease incidence was scored by a blinded veterinary pathologist using serial H&E and Picrosirius Red stained slides. Ordinal scoring was defined by percent of total parenchyma affected on the sampled section: 0 = 0% of total parenchyma, 1 = < 5%; 2 = 6-10%; 3 = 11-50%; 4 = 51-95%; 5 = > 95%. Instances of rare and isolated alveolar septa with gentle fibrotic changes were excluded from scoring.

### Chemokine & Cytokine analysis

Chemokine and cytokine profiles of serum and lung samples were assessed using Immune Monitoring 48-plex mouse ProcartaPlex Panel kits (Invitrogen). Briefly, 50 μL of either a 1:4 dilution of serum or 50 μL straight clarified lung homogenate were incubated with magnetic capture beads containing analyte specific antibodies. After washing, 96-well plates containing samples and magnetic beads were incubated with detection antibodies and SA-PE. Results were collected using a MAGPIX machine (Luminex) and quantification was achieved by comparing to a standard curve; both were done in xPONENT software. Values below limit of detection (LOD) were set to LOD and hierarchical clustering heatmaps were generated with the Bioconductor R package, *ComplexHeatmap*, after scaling the values across samples.

### Preparation of lung cell suspensions for flow cytometric analysis

Enzymatic digestion of lung tissue was performed by intratracheal instillation via a 20-gauge catheter of 1 mL of 5 mg/mL collagenase I (Worthington Biochemical Corp, Lakewood, NJ) and 0.25 mg/mL DNase I (Sigma) prepared in RPMI media (Life Technologies, Carlsbad, CA) prior to instilling 0.5 mL of 1% (wt/vol) low melting agarose (Amresco, Solon, OH), similar to previous protocols (*78*). Lung were then incubated at 37°C for 30 minutes. Lung were then minced and triturated through a 5 mL syringe. Cell suspensions were then filtered through a 50 mL conical 100 μM filter (ThermoFisher, Pittsburgh, PA) before RBC lysis and stained as previously described.

### Multi-color flow cytometry

The prepared lung cells were suspended in approximately 1 mL of PBS buffer supplemented with 1.5 % (w/v) bovine serum albumin (Sigma) and 2 mM EDTA (Sigma). The total cell count determined by hemocytometer with trypan blue (VWR). For each sample 1.5 x 10^6^ cells first underwent Fc receptor blockade with rat anti-mouse Fc*γ*RIII/II receptor (CD16/32; BD Biosciences). After Fc receptor blocking for 5 minutes on ice, cells were surface stained using antibodies listed in Key Resources Table and as previously described (*78*). For intracellular staining, the cells underwent fixation and permeabilization with the Foxp3/Transcription Factor Staining Buffer Set (eBioscience, San Diego, CA). Fixed and permeabilized single cells suspensions were subsequently stained with intracellular antibodies (**Supplemental Table 8**) to characterize differences in specific populations.

The neutrophils and macrophage subpopulations were identified through gating, as demonstrated in prior reports and adapted from previously published methods (*80*).

Flow cytometry was performed using a Cytoflex flow cytometer (Beckman Coulter, Brea, CA) and analyzed using CytExpert (Beckman Coulter) software. To determine the total number of a specific population in the lung, we first calculated the population’s percentage with respect to the total live single cell population. Next, we multiplied this percentage to the total cell count as determined by hemocytometer measurements to calculate the specific population’s total number per mouse lung.

### Specimen Computed Tomography (CT) Imaging

Phosphotungstic acid (PTA) staining was performed to increase soft tissue conspicuity for specimen computed tomography (CT) imaging. Lungs were inflated and fixed with 10% formalin at 20 cm H_2_O pressure for seven days. Samples were initially washed 3X in 70% EtOH in 50 ml non-reactive tubes prior to staining. Each lung was then immersed in 0.3% (w/v) Phosphotungstic acid hydrate (Sigma-Aldrich P4006) in 70% EtOH for seven days on an oscillating table. They were subsequently air dried prior to imaging.

Specimen CT scanning of the dried lungs was performed on a Sanco µCT 40 (ScanCo Medical AG, Switzerland. Imaging was performed at 70kVP at 114 µA current and 200 ms integration time. Images were reconstructed using a conebeam algorithm at 16 µm voxel size in a DICOM file format. Images were viewed with ImageJ.

### RNA *in situ* hybridization, Immunohistochemistry, and Quantification

For histopathological analyses on mouse lung tissue sections, left lung lobes were stored in 10% phosphate buffered formalin for at least 7 days before transferring out of the BSL for further processing. Histopathological scoring was performed after tissue samples were embedded with paraffin, sectioned, and stained. Immunohistochemistry (IHC) was performed on paraffin-embedded lung tissues that were sectioned at 5 microns. This IHC was carried out using the Leica Bond III Autostainer system. Slides were dewaxed in Bond Dewax solution (AR9222) and hydrated in Bond Wash solution (AR9590). Heat induced antigen retrieval was performed for 20 min at 100°C in Bond-Epitope Retrieval solution 2, pH-9.0 (AR9640). After pretreatment, slides were incubated with primary antibodies (see Key Resources Table) for 1h followed with Novolink Polymer (RE7260-K) secondary. Antibody detection with 3,3’-diaminobenzidine (DAB) was performed using the Bond Intense R detection system (DS9263). Stained slides were dehydrated and coverslipped with Cytoseal 60 (8310-4, Thermo Fisher Scientific).

RNA-ISH was performed on paraffin-embedded 5 μm tissue sections using the RNAscope Multiplex Fluorescent Assay v2 or RNAscope 2.5 HD Reagent Kit according to the manufacturer’s instructions (Advanced Cell Diagnostics). Briefly, tissue sections were deparaffinized with xylene and 100% ethanol twice for 5 min and 1 min, respectively, incubated with hydrogen peroxide for 10 min and in boiling Target Retrieval Reagent (Advanced Cell Diagnostics) for 15 min, and then incubated with Protease Plus (Advanced Cell Diagnostics) for 15 min at 40°C. Slides were hybridized with custom probes at 40°C for 2 h, and signals were amplified according to the manufacturer’s instructions.

Stained mouse tissue sections were scanned and digitized by using an Olympus VS200 slide scanner. Images were imported into Visiopharm Software^®^ (version 2020.09.0.8195) for quantification. Lung tissue and probe signals for targeted genes detected by RNA-ISH were quantified using a customized analysis protocol package to 1) detect lung tissue using a decision forest classifier, 2) detect the probe signal based on the intensity of the signal in the channel corresponding to the relevant probe. The same methodology was applied to quantify CD4^+^ and CD8^+^ cells identified by IHC. Positive signals for CD4^+^ cells were determined using contrast of red-blue channels at a determined threshold to exclude background, similarly, CD8^+^ cells were determined using contrast of green-blue channels. All slides were analysed under the same conditions. Results were expressed as the area of the probe relative to total lung tissue area.

Paraffin-embedded mouse and human tissue sections (5 μm) were used for fluorescent IHC staining. According to the previously described protocol (*81*) sections were baked at 60 °C for 2-4 hours followed by a deparaffinization step including xylene and graded ethanol. Antigen retrieval was achieved after rehydration by boiling slides in 0.1M sodium citrate at pH 6.0 in a microwave. Slides were allowed to cool down and rinsed with distilled water before quenching of endogenous peroxidase was performed with 0.5% hydrogen peroxide in methanol for 15 min. After a PBS wash, slides were blocked with 4% normal donkey serum for 60 min at room temperature followed by incubation with primary antibodies (diluted in 4% normal donkey serum in PBST) at 4 °C overnight. Isotype control (species-matched gamma globulin) was diluted in the same manner as the primary antibody. Slides were incubated for 60 min at room temperature with secondary antibodies after being washed in PBST. Reduction of background staining was achieved by utilization of Vector® TrueVIEW Autofluorescence Quenching Kit (Vector laboratories). Tissue sections were covered in glass coverslips by adding ProLong Gold Antifade Reagent with DAPI (Invitrogen). Stained human tissue sections were scanned and digitized by using an Olympus VS200 slide scanner.

### GeoMx Digital Spatial Profiling

Five µm-thick FFPE sections were prepared using the RNAscope & DSP combined slide prep protocol from NanoString Technologies. Prior to imaging, mouse tissue morphology was visualized by IHC for CD45 and RNAscope for SARS-CoV-2 RNA, and DNA was visualized with 500 nM Syto83. Human tissue morphology was visualized by IHC for immune cell marker CD45/epithelial cell marker panCK/Syto83 and for KRT5 (IHC)/SARS-CoV-2 (RNA) on serial sections. Mouse or Human Whole Transcriptome Atlas probes targeting over 19,000 targets were hybridized, and slides were washed twice in fresh 2X SSC then loaded on the GeoMx Digital Spatial Profiler (DSP). In brief, entire slides were imaged at 20X magnification and 6-10 regions of interest (ROI) were selected per sample. ROIs were chosen based on serial hematoxylin and eosin-stained sections and morphology markers (mouse: DNA/CD45 IHC/SARS-CoV-2 RNA; human: CD45/PanCK/Syto83 IHC and SARS-CoV-2 RNA/KRT5/DAPI IHC on serial sections by a veterinary pathologist (S.A.M.). The GeoMx then exposed ROIs to 385 nm light (UV) releasing the indexing oligos and collecting them with a microcapillary. Indexing oligos were then deposited in a 96-well plate for subsequent processing. The indexing oligos were dried down overnight and resuspended in 10 μL of DEPC-treated water.

Sequencing libraries were generated by PCR from the photo-released indexing oligos and ROI-specific Illumina adapter sequences and unique i5 and i7 sample indices were added. Each PCR reaction used 4 μL of indexing oligos, 4 μL of indexing primer mix, and 2 μL of NanoString 5X PCR Master Mix. Thermocycling conditions were 37°C for 30 min, 50°C for 10 min, 95°C for 3 min; 18 cycles of 95°C for 15 sec, 65°C for 1 min, 68°C for 30 sec; and 68°C 5 min. PCR reactions were pooled and purified twice using AMPure XP beads (Beckman Coulter, A63881) according to manufacturer’s protocol. Pooled libraries were sequenced at 2×27 base pairs and with the dual-indexing workflow on an Illumina NovaSeq.

### Analysis of mouse GeoMx transcriptomic data

For mouse samples, raw count, 3rd quartile (Q3) normalized count data of target genes from ROIs were provided by the vendor, which were used as input to downstream analyses (**Supplemental Table 3**). Mouse Q3 normalized data were used for principal component analysis (PCA) using the R package *ade4* and visualized using *factoextra* package. Raw count data were used for differential expression analysis using the Bioconductor R package, *variancePartition* (*82*), with transformation of raw counts by voom method (*83*). The dream function from *variancePartition* allows fitting of mixed-effect models to account for ROIs obtained from the same animal, and assay slides as random-effect factors. Differentially expressed genes (DEGs) were defined as genes that passed the filters of Benjamini-Hochberg adjusted p-value < 0.05, and absolute log2 fold-change > 1. Pre-ranked gene set enrichment analysis (GSEA) was performed using the Bioconductor R package, *fgsea* (*84*), with gene set collections obtained from Gene Ontology Biological Process (*85*), and Reactome pathways (*86*). Various gene lists of interests were curated manually from published literature, and human gene symbols from references were converted into homologous mouse genes using bioDBnet (https://biodbnet-abcc.ncifcrf.gov/). Plots and hierarchical clustering heatmaps were generated using the R package, *ggplot2* (*87*), and *ComplexHeatmap* (*88*).

For the human samples, WTA + COVID-19 spike-in gene targets were assayed. FASTQ data were first converted to digital counts conversion (DCC) format. Probe outlier tests were performed on each set of negative probes (one set of negative probes for the WTA panel and one set for the COVID-19 spike-in panel). Specifically, for a given negative probe pool, the geometric mean of all counts (across all probes and all samples) was computed. A probe was identified as a low count outlier if its probe-specific geometric mean divided by the grand geometric mean was less than the threshold of 0.1. From the remaining probes, the Rosner Test was used to detect local outliers on a sample-specific case using the R package EnvStats (*89*) with parameters *k* equal to 20% of the number of negative probes and *alpha* equal to 0.01. A negative probe was considered a global outlier if it was found to be a local outlier in more than 20% of samples and was discarded from downstream analysis. For each panel pool, the negative probe geometric mean and geometric standard deviation were computed. The sample-specific limit of quantification (LOQ) was estimated from these moments by multiplying the geometric mean by the geometric SD and then squaring that quantity. Gene targets in the COVID-19 spike-in, which contain multiple probes per target, were collapsed to a single floating point value using the geometric mean. Following outlier filtering, the sequencing saturation for each sample was computed as the one minus the number of deduplicated reads divided by the number of aligned reads. One sample yielded a sequencing saturation below the 0.67 cutoff (range of other samples: 85.9-96.8) and was removed. Additionally, one sample had an LOQ more than 2.7 standard deviations from the mean in the WTA panel and 4.2 standard deviations from the mean in the COVID-19 spike-in pool and was removed from the analysis. Filtering gene targets was also performed. If a gene target was below LOQ in more than 10% of samples, it was filtered out. Following the above probe, sample, and target filtering steps, the data matrix was normalized using the Q3 method (see above).

Preliminary analysis of the log2 transformed and scaled Q3 normalized data identified a putative batch effect between two runs as identified using the PCA in the R package FactoMineR. The following batch correction algorithm was used before downstream data analysis. We first ensured that the batching factor was not itself confounded with Group (Healthy or COVID-19) or Region (alveolar, bronchiolar, disorganized). This was done by creating a design matrix and checking for any linearly dependent terms using the core R package *stats* (*90*). No factors were correlated with Batch using a correlation threshold of 0.3. Batch correction was performed for each gene target by modeling its log2 Q3 expression (dependent variable) in a mixed effect model that included a random intercept for the fixed portion and Batch as a random effect with random intercept. Modeling was done in the R package *lme4*. For each model, the residuals of the model were extracted and converted back to the linear scale. These residuals were then multiplied by the model’s estimated intercept (also linear scale) to shift the values to an intensity similar to the original Q3 data. To evaluate how well the above approach removed the batch effect, we regressed the first 5 PC scores against Batch for both the Q3 as well as the batch corrected (BC) data using a series of ANOVAs. Of the five PC axes, only the first was associated with the batching factor (P < 4e-36; all others, P > 0.23) in the Q3 data. Following correction, no axes were associated with Batch (all P > 0.80).

### Histological scoring of human COVID-19 lung tissue

The H&E stained regions of interest (ROI) were scored by a pulmonary pathologist (S.G.) grading each section on a semi-quantitative scale between zero and three, with zero representing a normal human lung section and three representing the most severe histologic change encountered in clinical practice. The features scored in each ROI are: interstitial inflammation, airspace fibrin exudates (acute phase of lung injury), the fibroblastic/organizing-phase of lung injury and mature fibrosis. Human donor information can be found in **Supplementary Table 6**.

### Analysis of human GeoMx transcriptomic data

For human samples, raw count and Q3 + batch corrected count data of target genes from ROIs were provided by the vendor (**Supplemental Table 7**). Prior to downstream analysis, Q3 + batch corrected data were log_2_ normalized. Principal component analysis were performed on the top 1,000 highly variable genes on the log normalize data. Coexpression network analysis was performed on 11,556 expressed genes using Weighted Gene Coexpression Network Analysis (WGCNA) R package (*91*). Differential gene and network expression between groups were evaluated under a linear mixed model approach accounting for multiple ROIs per donor using R package *Ime4*. Statistical significance of the estimates were evaluated with R package *lmerTest* (*92*), using the Satterthwaite’s degrees of freedom method. Sets of differentially expressed genes were tested for overrepresentation of the genes in the databases (GO: Biological Process, GO: Molecular Function, GO: Cellular Components, KEGG, and Reactome) using R package *enrichR* (*93*). For each network, genes were selected based on the degree of correlation with the network eigengene. To cluster ROIs obtained from healthy and COVID-19 donors, hierarchical clustering was performed based on the 50 most correlated network genes from each of the 7 identified networks using ward.D2 agglomeration method. As a result, healthy ROIs were separated from COVID-19 ROIs and COVID-19 ROIs were segregated into three subtypes, including COVID1, COVID2, and COVID3. Various plots and heatmaps were generated using the R packages ggplot2 and heatmap3 (*94*).

### Human lung tissue and quantification of Sirius Red and smooth muscle actin signals

Control lungs were obtained from lung transplant donors without any history of pulmonary disease whose lungs were unsuitable for transplant due to size mismatch provided by the University of North Carolina (UNC) Tissue Procurement and Cell Culture Core (institutional review board (IRB)-approved protocol #03-1396). COVID-19 autopsy lung tissue sections were obtained from Drs. Ross. E. Zumwalt (University of New Mexico, Albuquerque, NM), Edana Stroberg (Office of the Chief Medical Examiner, Oklahoma City, OK), Alain Borczuk (Weill Cornell Medicine, New York, NY), and Leigh B. Thorne (University of North Carolina at Chapel Hill (UNC), Chapel Hill, NC). Human donor information can be found in **Supplementary Table 6**. Early- and late-phase specimens were defined as autopsy tissues obtained ≤ 20 and > 20 days post an onset of symptoms, respectively.

Stained areas of Sirius Red and SMA detected by IHC in the alveolar regions were quantitated using Fiji software. Alveolar regions were randomly selected and cropped from the field. Optimized threshold value was determined by adjusting the threshold accurately representing the original images. The optimized threshold values were applied to identify Sirius Red or SMA signals. The Sirius Red or SMA-stained areas were measured and normalized to alveolar areas.

### *In vivo* Drug Treatment

EIDD-2801 (Emory Institute of Drug Design) was dissolved in a solution of 2.5% cremaphor (Sigma-Aldrich), 10% PEG 400 (Fisher Chemical), and 87.5% Molecular biology grade water (HyClone) via bath sonication at 37°C for 10 minutes, as described previously (*53*). Drug solution was made at a concentration of 62.5 mg/mL fresh daily for a final dose of 250 mg/kg per mouse (500mg/kg BID). Mice were dosed via oral gavage with 100uL of vehicle or EIDD solution twice daily beginning at 12 hours post infection and were dosed every 12 hours until 120 hours post infection.

Nintedanib (MedChemExpress) suspension was made in Molecular Biology Grade Water (HyClone) with 1% Tween-80 (Sigma-Aldrich) fresh daily at a concentration of 15mg/mL for a final dose of 60mg/kg per mouse (*95, 96*). Mice were dosed once daily via oral gavage with either 100uL of Nintedanib suspension or vehicle starting at 7 days post infection until final harvest at either 15 or 30 days post infection. Mouse serum was harvested at indicated time points after nintedanib administration, inactivated for BSL3 removal with 0.05% Triton-X100 and heating at 56°C, and was analyzed using ultra high-performance liquid chromatography time-of-flight mass spectrometry (UHPC-TOF MS). Samples were prepared by precipitating protein with acetonitrile (Sigma-Aldrich) containing diazepam (Cerilliant) as an internal standard. The supernatant was separated using a Flexar FX-20 UHPLC system (Perkin Elmer) with a Kinetex C18 biphenyl column (2.6 um 50 x 3 mm Phenomenex) at 45°C with 98% MS-grade water (Sigma-Aldrich), 10 mM ammonium acetate (Hagn Scientific), and 98% methanol (Sigma-Aldrich) 0.1% formic acid (Hagn Scientific) gradient elution at a flow rate of 0.6 mL/min. The Perkin Elmer Axion2 TOF mass spectrometer operated in positive-ion electrospray ionization (ESI+) mode was used to detect accurate mass spectra of nintedanib at 540.2605 [M+H]+. The method was linear from 1 to 500 ng/mL with a lower limit of detection of 1 ng/mL. The results for nintedanib concentration in mouse sera for this study was in agreement with the serum concentrations reported previously (*58*).

### Quantification and statistical analysis

Wilcoxon rank-sum test was used to test the difference in CD4+ or CD8+ T cells **(Fig. S4C, D)**, as well as Sirius red- or SMA-stained areas **(Fig. S8C, D)**, identified by IHC between two groups. Flow cytometry data were analyzed by Wilcoxon rank-sum test (**Fig. S4E**) or ANOVA followed by Sidak’s multiple comparisons test **(Fig. S4F-H)**. The difference in DSP Q3 normalized counts for targeted genes in ROIs between each condition and time point was statistically tested using a linear mixed-effect model using the R package *Ime4* (*97*), with condition and time point as fixed effects and replicate mice as random-effect factors (**Fig. 4C, S5D-E**). Statistical significance was evaluated with the R lmerTest package(*92*), using the Satterthwarte’s degrees of freedom method. Multiple post-hoc comparisons of subgroups were performed using the R multcomp package (Hothorn T, 2008). P < 0.05 was considered statistically significant.

### Data and material availability

All relevant data is included in this article. SARS-CoV-2 MA10 is available from BEI resources. Reagents and resources available upon request to corresponding author and with material transfer agreement.

## List of Supplementary Materials

**Supplemental Fig. 1: Micro-CT scans of mouse lungs reveal pulmonary disease.**

**Supplemental Fig. 2: SARS-CoV-2 MA10 infection causes lung damage in young surviving mice.**

**Supplemental Fig. 3: SARS-CoV-2 MA10 induces local and systemic cytokine and chemokine responses.**

**Supplemental Fig. 4: SARS-CoV-2 MA10 induces robust immune cell infiltration.**

**Supplemental Fig. 5: Upregulation of ISG expression after SARS-CoV-2 infection.**

**Supplemental Fig. 6: SARS-CoV-2 MA10 infection causes transient loss of club and ATII cells.**

**Supplemental Fig. 7: Dynamics of ADI/DATP/PATS cell fates.**

**Supplemental Fig. 8: SARS-CoV-2 MA10 pathogenesis closely resembles late human COVID-19 disease.**

**Supplemental Table 1: Cytokine and chemokine protein levels in SARS-CoV-2 MA10 infected young and old mice**

**Supplemental Table 2: Gene lists in heat maps**

**Supplemental Table 3: Mouse whole transcriptome GeoMx data**

**Supplemental Table 4: Mouse GeoMx differential gene expression analysis**

**Supplemental Table 5: Mouse GeoMx pathway enrichment analysis**

**Supplemental Table 6: Human donor demographics**

**Supplemental Table 7: Human whole transcriptome GeoMx data**

**Supplemental Table 8: Reagent and Resource descriptions**

**References and Notes:**

## Acknowledgements

This project was funded in part by the National Institutes of Health, Department of Health and Human Service awards: National Institute of Allergy and Infectious Diseases (AI110700, AI108197, AI142759 and U54 CA260543 to RSB, T32 AI007419 to KHD and EJF), an animal models contract from the NIH (HHSN272210700036I 75N93020F00001 to RSB), National Institute of Diabetes and Digestive and Kidney Diseases (P30 DK065988 to RCB), National Heart, Lung, and Blood Institute (R01 HL152077 and K08 HL129075 to JRM; Cystic Fibrosis Foundation (RDP BOUCHE15R0 to RCB and BOUCHE19XX0 to RCB and KO, and OKUDA20G0 to KO). North Carolina Policy Collaboratory at the University of North Carolina at Chapel Hill with funding from the North Carolina Coronavirus Relief Fund established and appropriated by the North Carolina General Assembly. The authors appreciate Yasue Kato and Ella R. Strickler for image processing. Histopathology was performed by the Pathology Services Core at the University of North Carolina at Chapel Hill, which is supported in part by an NCI Center Core Support Grant (5P30CA016080-42).

KHD, SRL, RSB are listed on a pending patent for the SARS-CoV-2 MA10 virus.

AV, TH, ML, SJP, YL are employees and shareholders of NanoString Technologies, Inc.

