## Supplemental Figures for "A model of persistent post SARS-CoV-2 induced lung disease for target identification and testing of therapeutic strategies"

Fig. S1

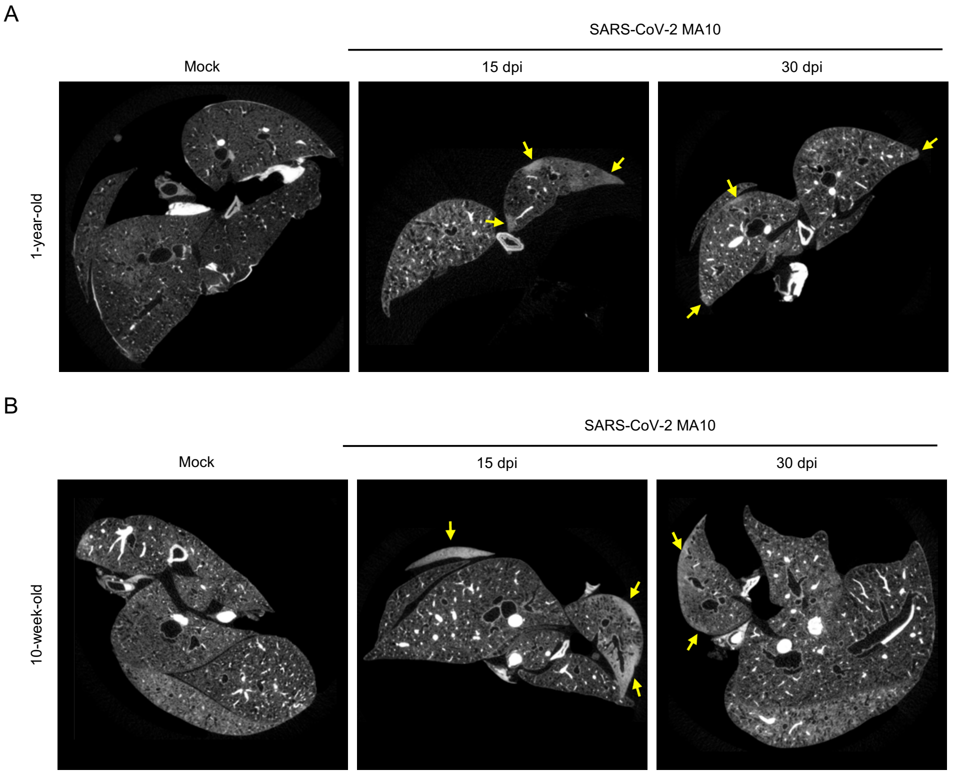


**Supplemental Fig. 1: Micro-CT scans of mouse lungs reveal pulmonary disease.** Representative images of specimen micro-CT scans from **(A)** 1-year-old mice and **(B)** 10-week-old mock and SARS-CoV-2 MA10 infected mice at 15 and 30 dpi. Arrows indicate areas of dense ground glass opacity and consolidation representing peripheral fibrosis

Fig. S2

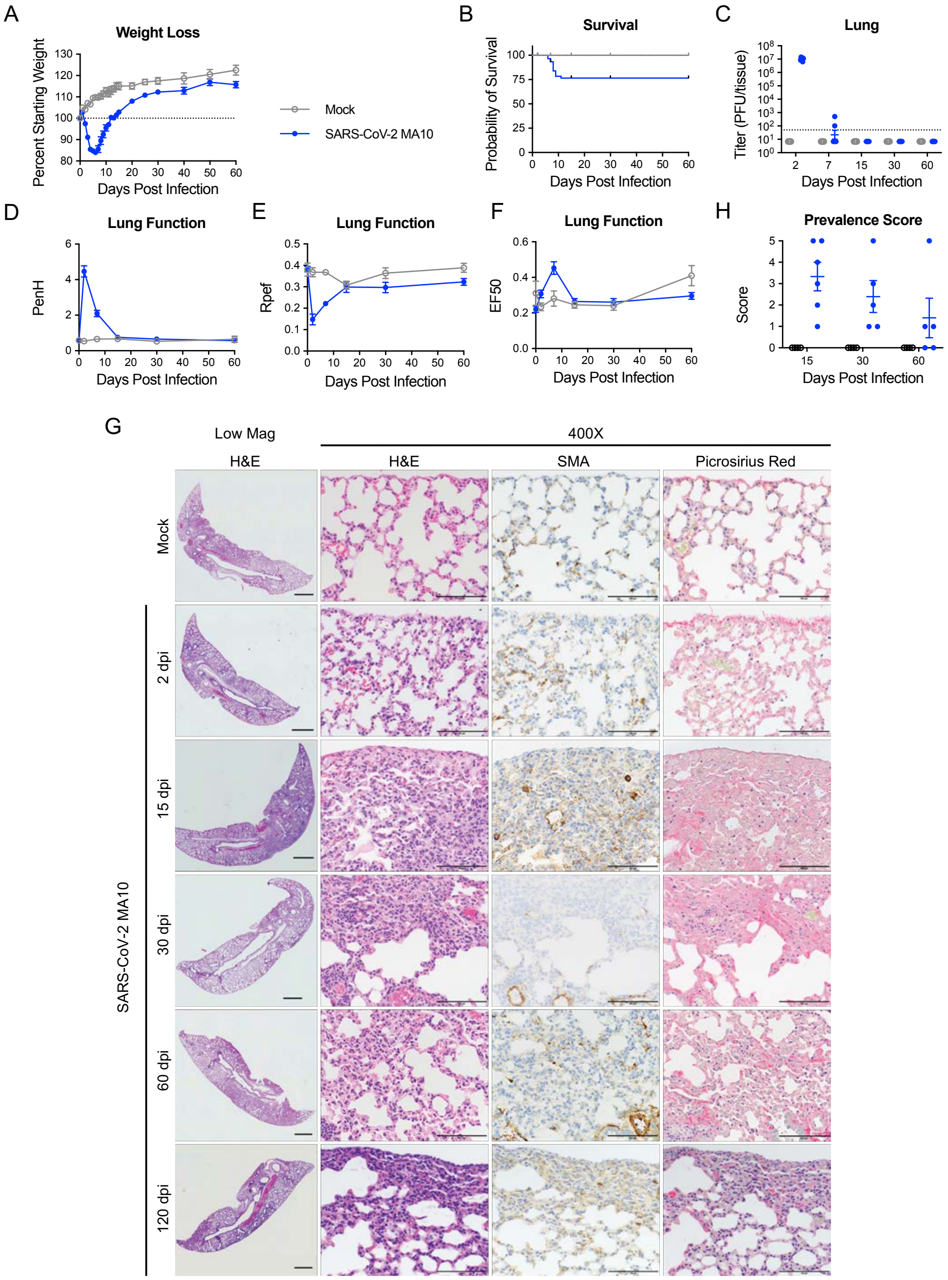


**Supplemental Fig. 2: SARS-CoV-2 MA10 infection causes lung damage in young surviving mice.** 10-week-old female BALB/c mice were infected with $10^4$ PFU SARS-CoV-2 (n=66) MA10 or PBS (n=24) and monitored for **(A)** percent starting weight and **(B)** survival. **(C)** Log transformed infectious virus lung titers were assayed at indicated time points. Dotted line represents limit of detection. Undetected samples are plotted at half the limit of detection. **(D-F)** Lung function was assessed by whole body plethysmography for **(D)** PenH, **(E)** Rpef, and **(F)** EF50. **(G)** Histopathological analysis of lungs at indicated time points. H&E: hematoxylin and eosin. SMA: immunohistochemistry for smooth muscle actin. Picrosirius Red directly stains collagen fibers. Image scale bars represents 1000 µm for low magnification and 100 µm for 400X images. **(H)** Disease incidence scoring at indicated time points: 0 = normal; 0 = 0% of total parenchyma, 1 = < 5%; 2 = 6-10%; 3 = 11-50%; 4 = 51-95%; 5 = > 95%. Graphs represent individuals necropsied at each timepoint (C, H), with the average value for each treatment and error bars representing standard error of the mean, calculated in Prism 9 (A-H). Mock infected animals represented by open gray circles and SARS-CoV-2 MA10 infected animals are represented by closed blue circles.

Fig. S3

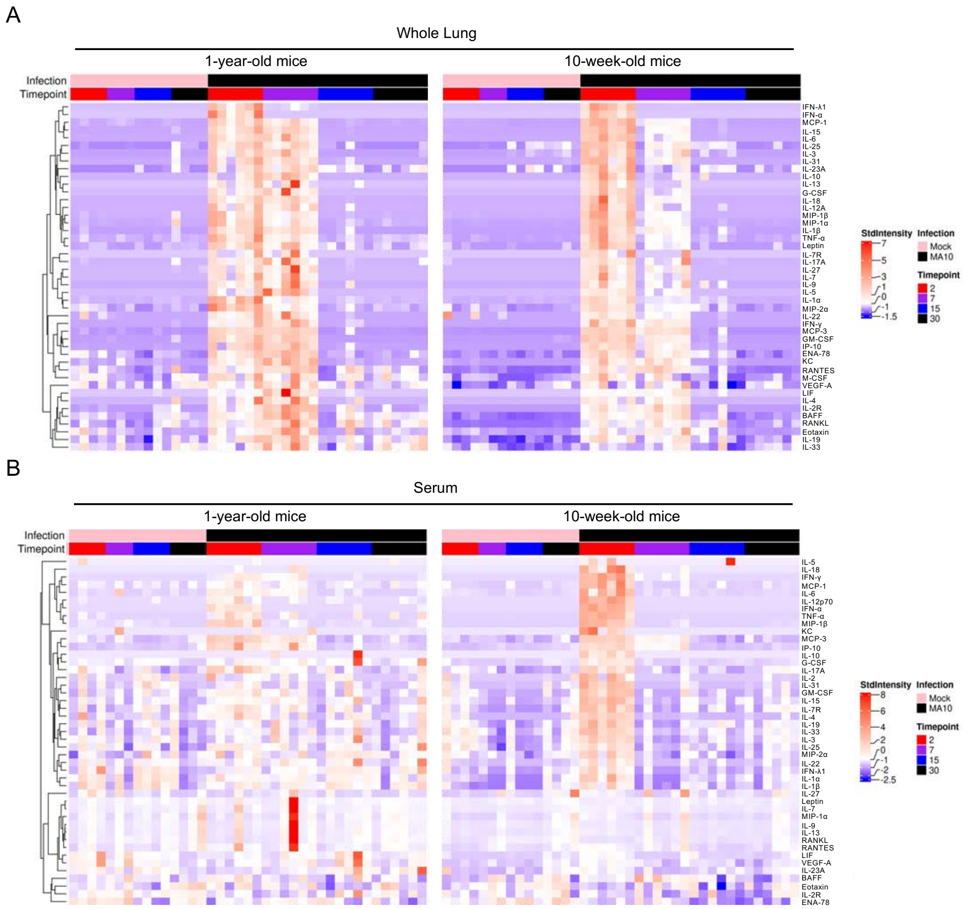


**Supplemental Fig. 3: SARS-CoV-2 MA10 induces local and systemic cytokine and chemokine responses.** Heatmaps of cytokine and chemokine protein levels in lung tissue homogenate **(A)** or serum **(B)** in 1-year-old or 10-week-old mock or SARS-CoV-2 MA10 infected mice at 2, 7, 15, and 30 dpi.

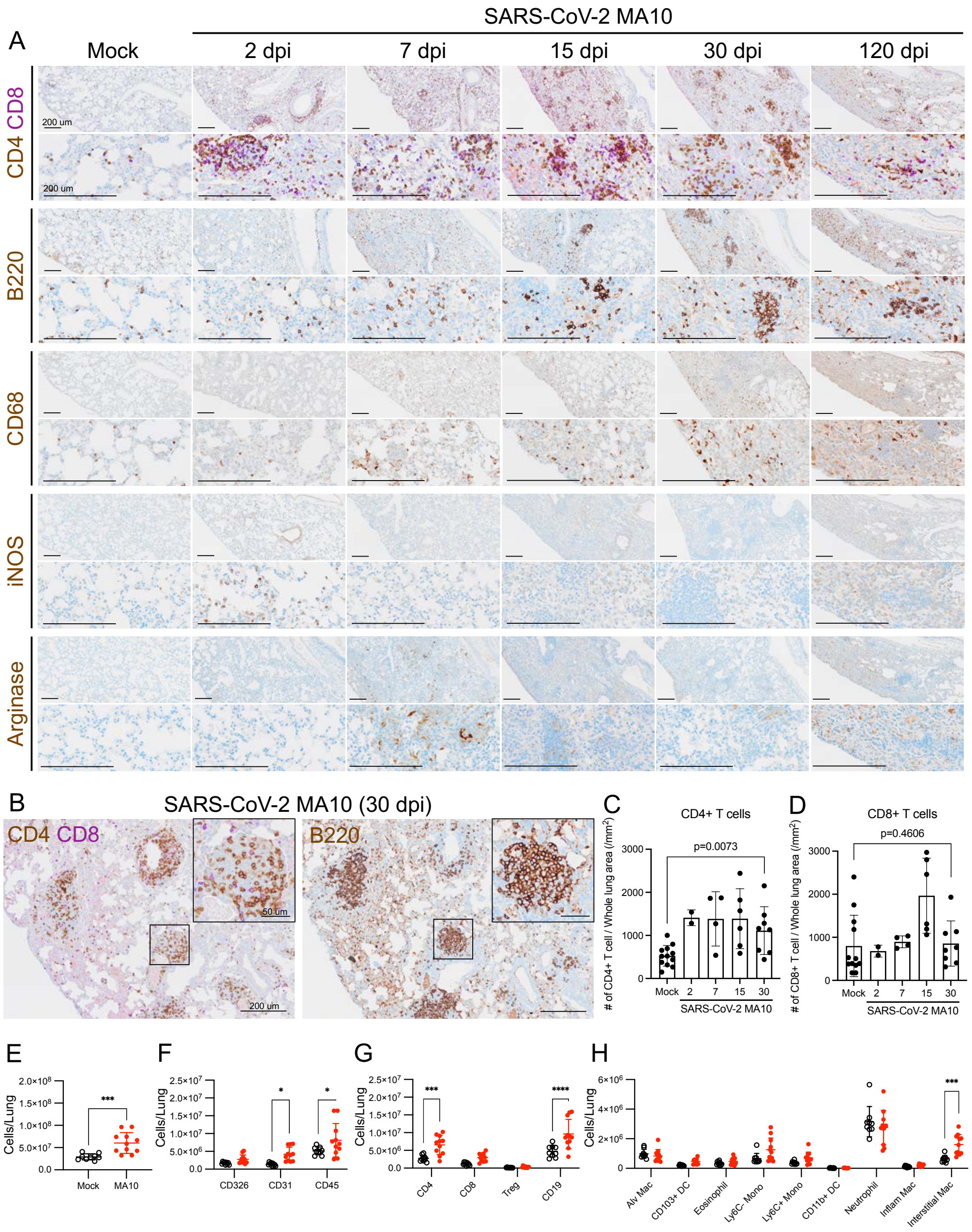


**Supplemental Fig. 4: SARS-CoV-2 MA10 induces robust immune cell infiltration. (A)** Immunohistochemistry of $CD4^{+}$ and $CD8^{+}$ T cells, $B220^{+}$ B cells, $CD68^{+}$ macrophages, and $iNOS^{+}$ M1, and $Arginase^{+}$ M2 macrophages at indicated time points after infection compared to mock. Scale Bars = 200 µm. **(B)** Immunohistochemistry depicting alveolar lymphocyte aggregates consisting of $CD4^{+}$, $CD8^{+}$, and $B220^{+}$ B cells at 30 dpi. Scale Bars = 200 µm (low power) and 50 µm (insets). **(C-D)** Quantification of **(C)**

CD4$^+$ and **(D)** CD8$^+$ T cells based on immunohistochemistry **(A)**. Graphs represent individuals necropsied at each timepoint with the average value for each treatment and error bars representing standard error of the mean. Wilcoxon rank-sum test was used to test the difference in CD4$^+$ or CD8$^+$ T cells identified by immunohistochemistry between two groups **(C,D)**. **(E-H)** Quantification of cells by flowcytometry: **(E)** total live cell counts; **(F)** total CD326$^+$ epithelial, CD31$^+$ endothelial, and CD45$^+$ immune cells; **(G)** CD4$^+$, CD8$^+$, regulatory T cells (Tregs), and CD19$^+$ B cells; **(H)** subsets of myeloid lineage immune cells. Graphs include individuals selected for flow cytometry with average and error bars representing the standard error of the mean, mock individuals are represented by open black circles and SARS-CoV-2 infected are represented by closed red circles (**E-H**). Flow cytometry data analyzed by Wilcoxon rank-sum test (**E**) or ANOVA followed by Sidak's multiple comparisons test (**F-H**).

Fig. S5

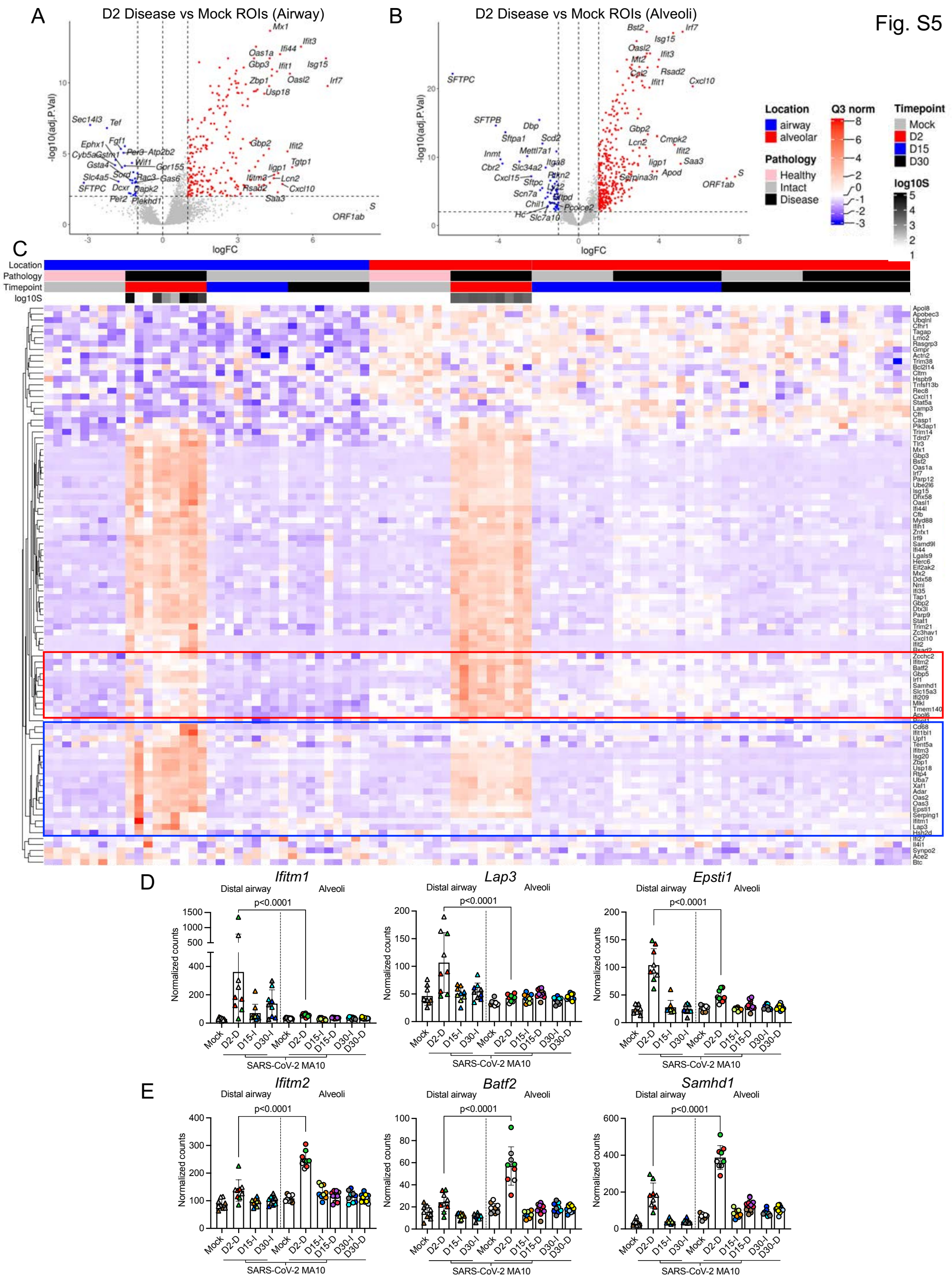


**Supplemental Fig. 5: Upregulation of ISG expression after SARS-CoV-2 infection. (A-B)** Volcano plots of DSP differentially expressed genes (DEGs) in SARS-CoV-2 MA10 infected **(A)** distal airway or **(B)** alveolar ROIs at 2 dpi *vs.* mock. **(C)** DSP heatmap of selected interferon stimulated genes (ISGs) in alveolar and distal airway epithelial ROIs across all time points. Red square highlights ISGs highly expressed in alveolar disease ROIs compared to distal airway disease ROIs at 2 dpi. Blue square highlights

ISGs highly expressed in distal airway disease ROIs compared to alveolar disease ROIs at 2 dpi. **(D-E)** DSP Q3 normalized counts for selected ISGs highly upregulated in **(D)** distal airway disease ROIs or **(E)** alveolar disease ROIs at 2 dpi. Graphs represent all ROIs selected with each unique color and symbol representing one animal, bars represent average value of each group **(D-E)**. The difference in DSP Q3 normalized counts for targeted genes in ROIs between each condition and time point was statistically tested using a linear mixed-effect model with condition and time point as fixed effects and replicate mice as random-effect factors.

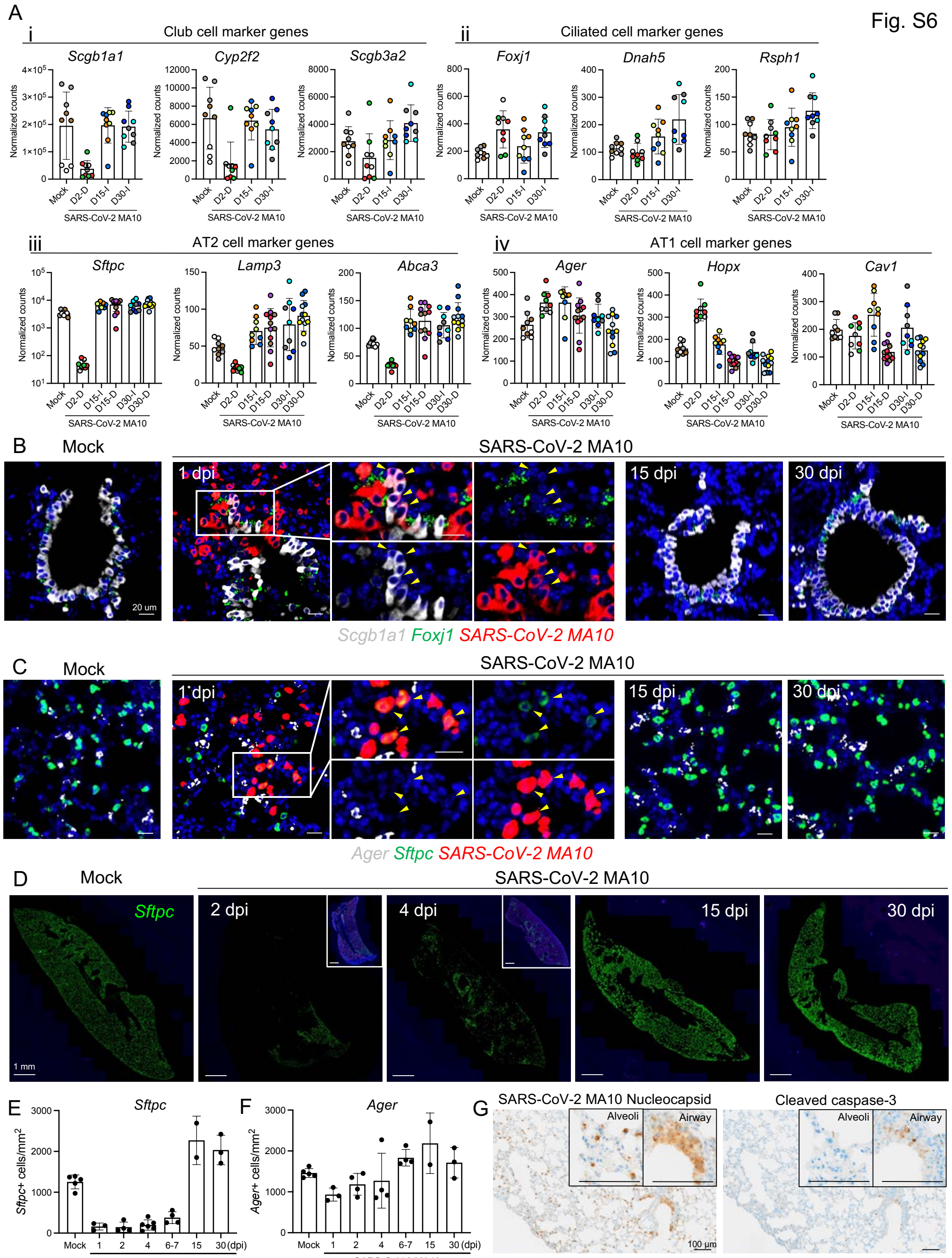


**Supplemental Fig. 6: SARS-CoV-2 MA10 infection causes transient loss of club and ATII cells. (A)** DSP Q3 normalized counts for club (i), ciliated (ii), AT2 (iii), and AT1 (iv) cell markers in mock, infected diseased (D), or intact (I) distal airway (i, ii) or alveolar (iii, iv) ROIs. Graphs represent all ROIs selected with each unique color and symbol representing one animal, bars represent average value of each group with error bars representing standard error of the mean. **(B)** RNA-ISH for club (*Scgb1a1*) and ciliated

(*FoxJ1*) markers at indicated timepoints. Arrow heads indicate colocalization of *SARS-CoV-MA10 RNA* with *Scgb1a1*. **(C)** RNA-ISH for AT2 (*Sftpc*) and AT1 (*Ager*) markers at indicated timepoints. Arrow heads indicate colocalization of *SARS-CoV-MA10 RNA* with *Sftpc*. **B-C**: Scale Bars = 20 μm. **(D)** Low magnification RNA-ISH for *Sftpc* at indicated time points. Scale Bars = 1 mm. **(E-F)** Quantification of RNA ISH for **(E)** *Sftpc* or **(F)** *Ager* at indicated timepoints with average and standard error of the mean plotted. **(G)** Immunohistochemistry for SARS-CoV-2 N and cleaved caspase-3 at 2 dpi. Scale Bars = 100 μm.

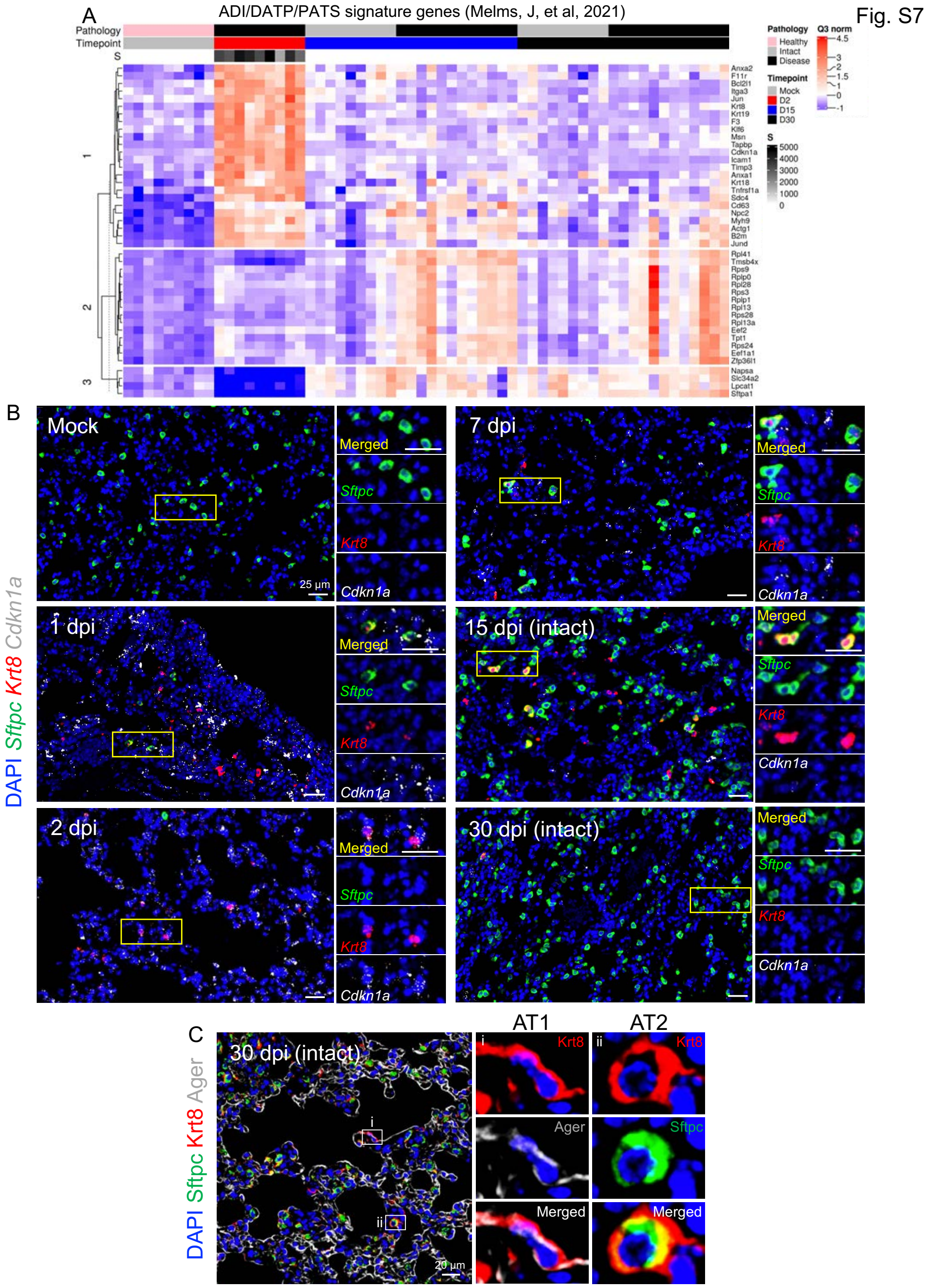


**Supplemental Fig. 7: Dynamics of ADI/DATP/PATS cell fates. (A)** DSP heatmap of ADI/DATP/PATS signature genes identified in COVID-19 autopsy lungs(*45*). **(B)** RNA-ISH of *Sftpc* and ADI/DATP/PATS cell markers *Krt8* and *Cdkn1a* over time course after SARS-CoV-2 MA10 infection with selected areas of interest. **(C)** Immunohistochemistry of Krt8 with AT1 (Ager) (i) and AT2 (Sftpc) (ii) cell markers.

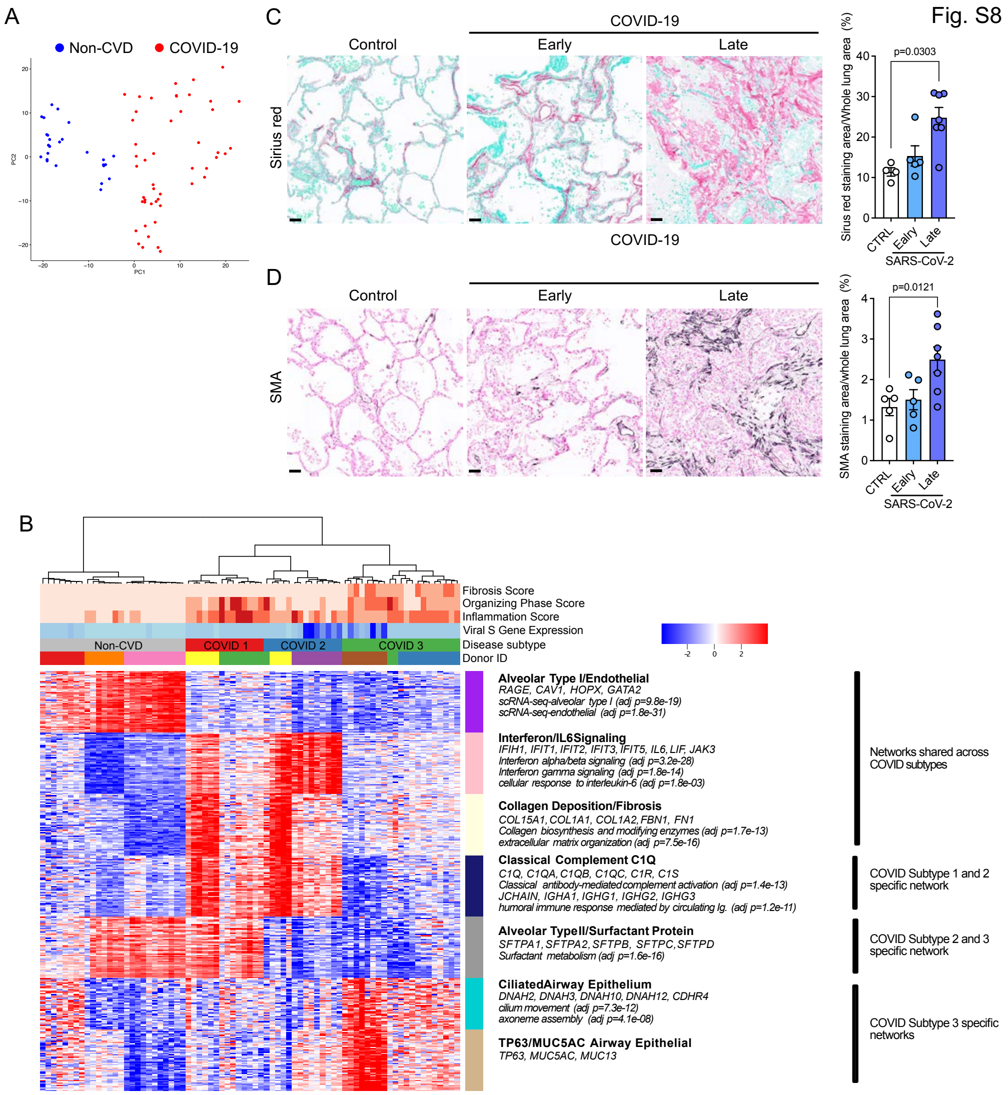


**Supplemental Fig. 8: SARS-CoV-2 MA10 pathogenesis closely resembles late human COVID-19 disease. (A)** PCA plot of DSP alveolar ROIs selected in non-COVID (non-CVD) (n=3) and COVID-19 (n=5) human donor lungs. **(B)** DSP heatmap of 7 identified networks in alveolar ROIs obtained from non-COVID control and COVID-19 subjects. Hierarchical clustering segregated COVID-19 ROIs into three subtypes (COVID 1, COVID 2, and COVID 3). DSP Q3 normalized counts of SARS-CoV-2 Spike (*S*) gene and histopathological scoring of alveolar ROIs for inflammation, organizing phase of lung injury, and fibrosis are shown. **(C-D)** Histopathological analysis of fibrotic features in control (n=4), early (n=5), and late (n=6) COVID-19 autopsy lungs by Sirius red for collagen deposition (**C**) and immunohistochemistry for smooth muscle actin (SMA) (**D**). Scale Bars = 50 μm. Sirius red and SMA immunohistochemistry quantification represents average value for each group with error bars representing standard error of the mean. Wilcoxon rank-sum test was used to test the difference in Sirius red- or SMA-stained areas identified by immunohistochemistry between two groups.
